# All-*trans* retinoic acid and fluid transport in myopigenesis

**DOI:** 10.1101/2025.02.05.636685

**Authors:** Mariia Dvoriashyna, Melissa Bentley-Ford, Jianshi Yu, Saptarshi Chatterjee, Machelle T. Pardue, Maureen A. Kane, Rodolfo Repetto, C. Ross Ethier

**Affiliations:** School of Mathematics and Maxwell Institute for Mathematical Sciences, University of Edinburgh, Edinburgh, UK; Department of Ophthalmology, Emory University School of Medicine, Atlanta, GA, USA; Center for Visual and Neurocognitive Rehabilitation, Atlanta VA Healthcare System, Atlanta, GA, USA; Department of Pharmaceutical Sciences, University of Maryland, Baltimore, MD, 21201, USA; Department of Civil, Chemical and Environmental Engineering, University of Genoa, Genoa, Italy; Wallace H. Coulter Dept. of Biomedical Engineering, Georgia Institute of Technology & Emory University, Atlanta, GA, 30332, USA

**Keywords:** myopia, all-trans retinoic acid, mathematical modelling, unconventional outflow

## Abstract

Myopia, or near-sightedness, is rapidly growing in prevalence, with significant long-term implications for ocular health. There is thus great impetus to better understand molecular signaling pathways leading to myopia. We and others have reported that all-trans retinoic acid (atRA) is involved in myopigenic signaling, yet the understanding of how atRA is transported and exerts a myopigenic influence is poor. Here we measured the concentrations of atRA in the serum in wild-type C57BL/6 mice under control conditions and after atRA feeding, previously shown to induce myopia. We also developed a mathematical model that describes fluid fluxes and the advective-diffusive transport of atRA in choroid and sclera, including atRA synthesis in the choriocapillaris, atRA degradation by scleral cells, and binding of atRA to the carrier protein serum albumin. This model, developed for both mice and humans, showed that atRA produced in the choriocapillaris was able to permeate well into the sclera in both mice and humans at biologically-relevant concentrations, and that atRA feeding greatly increased tissue levels of atRA across both the choroid and sclera. We were also able to identify which parameters most influence atRA concentration in ocular tissues, guiding future experimental work. Our findings support atRA’s role in myopigenic signaling.

## 1. Introduction

Myopia, or near-sightedness, is a large and growing public health problem, with 50% of all people predicted to be myopic by the year 2050 [1]. Because myopia is a risk factor for several potentially blinding ocular diseases [2,3], the increased incidence of myopia has concerning consequences for vision, both directly and indirectly. There is thus significant impetus to develop better treatments for myopia, yet this development is hindered by a weak understanding of myopigenesis.

In almost all cases of human myopia, the sclera grows excessively, leading to an elongated eye [4]. During development, maturation and myopigenesis, scleral growth is mediated by a retinoscleral signaling pathway [5,6], motivating interest in understanding the mechanisms underlying this pathway. Our group and others [7–12] have identified all-*trans* retinoic acid (atRA) as a likely key player in this signaling pathway: notably, in mice, exogenous delivery of atRA by feeding causes myopia that is remarkably similar to that seen in classic models of myopigenesis, such as lens defocus [7].

atRA is a naturally occurring derivative of vitamin A that plays a key role in signaling during development, where it activates nuclear retinoic-acid receptors that modulate transcription [13,14]. It is highly hydrophobic [15] and thus is primarily transported cytoplasmically and in the extracellular space by binding to carrier proteins such as cellular retinoic acid-binding protein 2 (CRABP2) and serum albumin (SA) [16–18]. For example, in the blood, albumin was identified as an important carrier of atRA [19]. In the chick eye, apolipoprotein A-I was identified as a major retinoic acid-binding protein [20], although this has not been confirmed in mammalian eyes. Conversely, based on its relative abundance, we have suggested that SA may be the dominant atRA carrier in the extravascular space, even though the binding affinity of atRA to SA is only of the order of 0.1 μM^-1^ [21].

atRA can also be produced within relevant ocular tissues, as identified by the presence of cells expressing isoforms of retinaldehyde dehydrogenase, enzymes which irreversibly transform retinaldehyde into atRA. For example, several labs [9,22] identified the choroid as the main site of atRA generation during myopigenesis, although this finding was not supported in one study [8]. Further, retinaldehyde dehydrogenase Raldh2 was the only isoform detectable in the choroid [23], with Raldh2-positive cells localized around choroidal vessels in a distribution that apparently changes during myopigenesis [23].

In view of the complexity of the retinoscleral signaling pathway, it would be useful to have a better understanding of how atRA is transported in relevant ocular tissues (choroid, sclera). Visualization of atRA within tissue is difficult due to its low abundance, its weak native fluorescence and its complexation with carrier proteins, and thus we investigated this issue through computational modeling, informed by direct (bulk) measurements of atRA levels in relevant tissues before and after atRA feeding in mice. To model atRA transport, it is necessary to have a clear understanding of fluid fluxes within relevant tissues; thus, we also modeled fluid transport processes in the choroid and sclera. Our overall goal was to identify physical factors that significantly affected atRA transport and signaling in myopigenesis.

## 2. Methods

### 2.1 Overview

In general, transport processes within tissues are complex; the situation with atRA is doubly so because of its binding to carrier protein(s), which implies the need to track atRA+protein complex(es) plus the unbound protein(s). Here we lay out, at a high level and in point form, the main elements of our experimental approach and mathematical model, with details provided below.

- We experimentally measure plasma concentrations of atRA in two cohorts of wild-type mice: an experimental cohort that has received atRA feeding (exogenous delivery), and a control cohort that has not received atRA feeding.
- We create a mathematical model of atRA transport within the eye, with the following key features:
  - We consider a simplified “slice” domain allowing us to describe the influx, production, movement, and degradation of atRA in the interior-exterior direction within the eye (Figure 1). We restrict the domain to tissues exterior to the retinal pigment epithelium (RPE), i.e. to the choroid, suprachoroidal space (SCS), and sclera. This domain allows us to model the effects of oral atRA ingestion, since consumed atRA enters the blood (see below) and is assumed to be delivered to the eye primarily via leakage from the choroidal vasculature.
  - We account for production of atRA by cells resident with the choroid, and for degradation of atRA by CYP26 enzyme-containing cells within the sclera.
  - We allow atRA (plus a carrier protein) to be transported by diffusion and convection, which requires us to consider several sources of fluid convection: fluid pumping by the sPE, flow from the SCS across the sclera (uveoscleral outflow), and net flow from the SCS into choroidal vessels (uveovortex outflow) [24]. These fluid fluxes are coupled to a lumped parameter model of fluid transport in the eye (Goldmann’s equation [25]) so that the intraocular pressure (IOP) is computed for a given rate of inflow of aqueous humor.
  - We use one set of parameter values suitable for human eyes, and a second set suitable for mice. We have attempted to be parsimonious in the introduction of parameters, but inevitably, some quantities are introduced whose values are unknown. We determined these unknown values by an optimization process in which we compared model outputs with good-quality experimental measurements.
  - We assume steady state, since the time course of myopigenesis is days to weeks in experimental settings, while equilibration time scales for protein diffusion across the sclera are of order hours.

**Figure 1:**
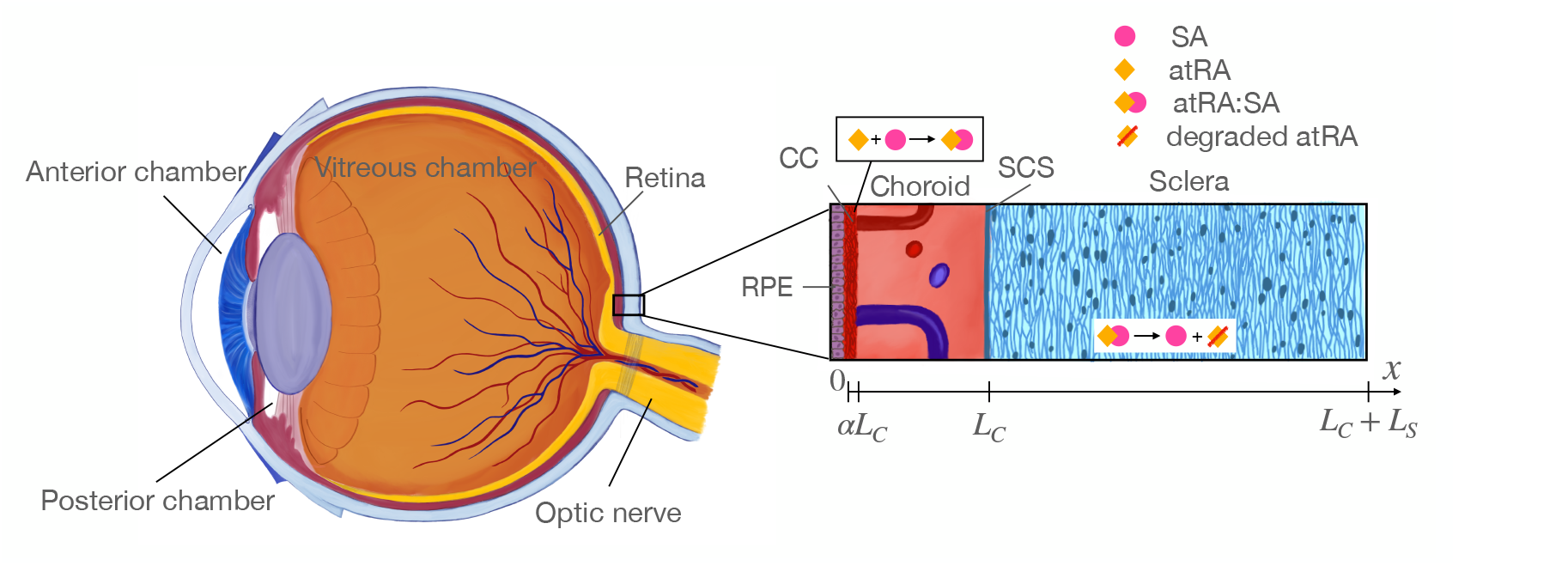
Overview of relevant ocular anatomy (left) and a schematic expanded view of the essential aspects of the mathematical model (right). Thicknesses are not to scale for the mouse eye. The right panel shows atRA entering the choroid (with local production and leakage from the choroidal vasculature lumped together) and binding to serum albumin (SA) to form an atRA:SA complex, which is then transported through the choroid into the sclera where it is taken up by scleral cells scleral cells to trigger signaling before atRA is eventually enzymatically degraded. Relevant lengths are shown below the image, in which *αL*_*C*_ is the fraction of the choroid occupied by the choriocapillaris, where atRA is synthesized and blood-extravascular exchange occurs.

### 2.2 Animal Husbandry and Sample Collection

All animal procedures adhered to the ARVO Statement for the Use of Animals in Ophthalmic and Vision Research, and were approved by the Institutional Animal Care and Use Committee at the Atlanta VA Healthcare System or Emory University School of Medicine. Male and female C57BL/6J mice (Jackson Laboratory, Bar Harbor, Maine, USA) were housed in the Emory animal facility on a 12-hour light-dark cycle. Mice approximately 4-5 weeks of age were fed with either 1% atRA in sugar pellets (experimental group 25 mg atRA/kg body weight) or sugar pellets only (control group).

At defined times after feeding, animals were anesthetized via intraperitoneal injection of ketamine (80-100mg/kg) and xylazine (5-10mg/kg) approximately 10 minutes prior to blood collection. Blood was collected in serum collection tubes (SARSDEDT, 41.1378.005; Newton NC). Samples were left at room temperature for 15-30 minutes followed by centrifugation at 2,000g for 10 minutes at 4°C. Serum was transferred to and stored in clean Eppendorf tubes. Samples were stored at −80 °C until they were shipped to the University of Maryland on dry ice for mass spectrometry.

We previously reported on the protocol for measurements of atRA levels in the RPE, choroid, and sclera [7], and use that existing data here. Note that we denote the RPE plus choroid plus sclera as “R/C/S”, which was treated as a single tissue source due to the difficulty of microdissecting the individual constituent tissues.

### 2.3 Mass Spectrometric Measurements of atRA Levels

Quantification of atRA in serum was performed as previously described [7,26–28]. All handling of retinoids and extraction of retinoid-containing fluid was performed under yellow light. In brief, volume of serum was determined, typically between 100-200 µL. Extraction of atRA from serum was accomplished by a two-step liquid-liquid extraction, using 4,4-dimethyl-RA as an internal standard, as previously described in detail [28]. Liquid chromatography-multistage-tandem mass spectrometry was used to quantify atRA, according to a previously described and validated method capable of distinguishing atRA from other biologically occurring isomers (9-cis-RA, 9,13-di-cis-RA, and 13-cis-RA) [26]. Liquid chromatography-multistage-tandem mass spectrometry was performed on a Prominence UFLC XR liquid chromatography system (Shimadzu, Columbia, MD), coupled to a 6500+ QTRAP hybrid tandem quadrupole mass spectrometer (AB Sciex, Foster City, CA), using atmospheric pressure chemical ionization operated in positive ion mode.

### 2.4 Mathematical Model of atRA Transport

We consider a domain *x* = [0, *L*_*C*_ + *L*_*S*_], where *L*_*C*_ is the thickness of the choroid and *L*_*S*_ is the thickness of the sclera (Figure 1). Note that this approach neglects antero-posterior and azimuthal variations in transport, and thus the domain should be considered as being located at an “average” location within the eye. As shown in Figure 1, the *x*-axis is normal to the tangent plane to the choroidal surface, has origin at the RPE/choroidal boundary, and points exteriorly.

#### 2.4.1 Fluid transport

Aqueous humor can exit the eye via two pathways: the so-called unconventional and conventional drainage pathways. We enforce the coupling of the conventional and unconventional pathways via the Gold-mann equation

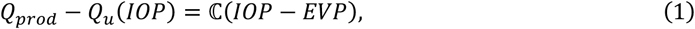

where *Q*_*prod*_ is the rate of production of aqueous humor by the ciliary processes, *Q*_*u*_(*IOP*) is the rate of aqueous outflow through the unconventional pathway, 𝕔 is aqueous outflow facility, *IOP* is (unknown) intraocular pressure, and *EVP* is episcleral venous pressure. Here 𝕔 and *EVP* are taken to be constant. Note that in (1) we have explicitly indicated that the unconventional outflow rate can depend (in part) on *IOP*.

We treat the choroid and sclera as porous media within which the flow is described by Darcy’s law

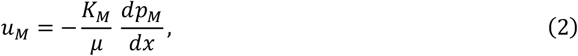

where *u*_*M*_, *K*_*M*_, *µ* and *p*_*M*_ are the superficial velocity, Darcy permeability, fluid viscosity and pressure in medium *M* (*M* = *C* or *S*, denoting choroid and sclera, respectively). We couple Darcy’s law with conservation of fluid mass, which in the 1D incompressible case takes the form

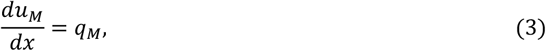

where *q*_*M*_ is a source term (volumetric rate of fluid entering the tissue per unit tissue volume). Within the sclera we assume there are no fluid sources, so that *q*_*S*_ = 0, and by assuming that *K*_*S*_ is constant, we can simply write

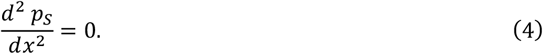

The situation in the choroid is more complex, since fluid can be exchanged between the choroidal vessels and the extravascular space by transmural filtration, i.e. *q*_*C*_ ≠ 0. We describe this process using Starling’s law

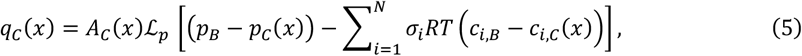

where *A*_*C*_(*x*) is the surface area of choroidal capillaries per unit tissue volume, ℒ_*p*_ is the mural hydraulic conductivity of these capillaries, *σ*_*i*_ is the reflection coefficient of the capillary walls to osmotically active component *i, R* is the universal gas constant, *T* is temperature, *c*_*i*_ is the concentration of osmotically-active component *i*, the subscript “B” refers to the blood plasma, and the summation is carried out over all *N* relevant osmotically-active components. Note that we have adopted the convention that *q*_*C*_ > 0 means a net fluid loss from filtering capillaries into the extravascular choroidal tissue. For purposes of this model, we consider a single “pool” of choroidal blood representing all blood within the choriocapillaris, and assume that this pool can be described by a single (average) pressure *p*_*B*_ and that the osmotically active solutes in the plasma can similarly be represented by a set of (average) concentrations *c*_*i,B*_. Because of the Starling resistor effect, the pressure in the vortex veins draining the choroid cannot drop below *IOP*. We therefore express the blood pressure in terms of the *IOP* plus a constant offset, i.e.

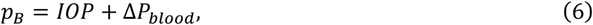

where Δ*P*_*blood*_ is an input parameter. Similarly, we assume there is an offset between *IOP* and the pressure in the SCS, *p*(*x* = *L*_*C*_), given by the expression

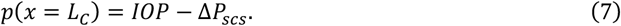

The parameter Δ*P*_*scs*_ is user-defined and is motivated by experimental measurements of Emi et al., who directly cannulated the suprachoroidal space in rabbits and showed an offset between *IOP* and SCS pressure over a range of IOPs [29].

The distribution of filtering capillaries across the thickness of the choroid is highly spatially non-uniform, i.e. capillaries are located within the choriocapillaris, which is adjacent to the RPE. We describe the spatial distribution of filtering capillaries using an empirical function *a*(*x*), which is maximum at the choroidal-RPE interface and decreases to zero with position across the choroid according to

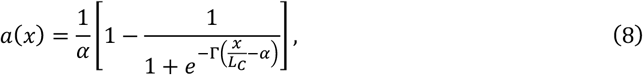

where Γ is a large user-defined constant (Γ ≫ 1) and *α L*_*C*_ is the characteristic length scale over which *a*(*x*) decreases, i.e. *a*(*x* = *αL*_*C*_) = 1/2*α*. In this work we used Γ = 100. Note that *a*(*x*) is a type of smoothed step function, and is normalized such that 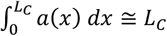, which allows us to write *A*_*C*_(*x*) in equation (5) as 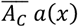, where 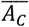 is the average value of *A*_*C*_(*x*) in the choroid. This allows us to write

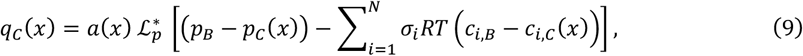

where we defined a modified permeability 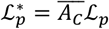, which is treated as a fitting parameter. Note that 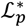 has units of 1/(s·Pa). Thus, the governing equation for the pressure within the choroid is obtained by combining (2), (3) and (9) to write

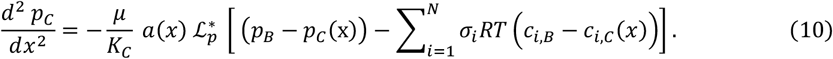

##### Boundary conditions for fluid transport

We account for fluid being pumped across the RPE into the choroid [30], assuming that the RPE pumps at a constant velocity *u*_*RPE*_ and hence imposing the condition:

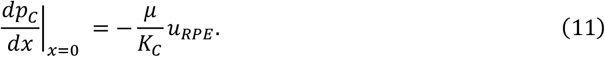

At the choroidal-scleral interface we require continuity of pressure and conservation of fluid mass

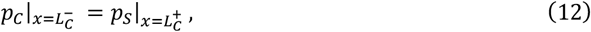

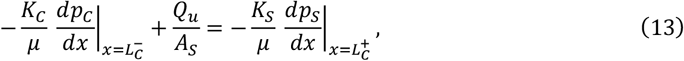

where *A*_*S*_ is the surface area of the choroidal-scleral interface and the term 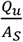 accounts for the volumetric flow rate of fluid entering the choroid and sclera via the suprachoroidal space due to unconventional outflow. *Q*_*u*_ is obtained from equation (1). Finally, the pressure at the external boundary of the sclera, *x* = *L*_*C*_ + *L*_*S*_, is set equal to orbital pressure

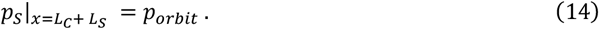

#### 2.4.2 Solute transport

We consider three species:

- Species 1: free atRA, which is required in our framework to account for synthesis of atRA by choroidal cells;
- Species 2: SA; and
- Species 3: atRA bound to SA, here denoted as atRA:SA.

The concentration of species *i* is denoted as *c*_*i*_ (*i* = 1,2,3). It is fairly simple to extend this framework to include other protein carriers of atRA, but as noted above, it seems likely that SA is an important atRA carrier in mouse and human eyes. Further, SA is the dominant determinant of the transmural osmotic pressure difference in the choriocapillaris and hence of transmural fluid transport; thus, we have here restricted attention to SA as the atRA protein carrier.

We describe species transport by the 1D steady convection-diffusion equation:

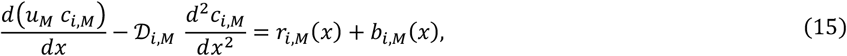

where 𝒟_*i,M*_ is the diffusivity of species *i* in tissue *M, r*_*i,M*_ is a source term due to chemical reactions (production and/or degradation of species *i*) and *b*_*i,M*_ is a source term due to transmural exchange of species *i* between the blood and the extravascular space.

Equation (15) evidently requires us to specify *r*_*i,M*(_ and *b*_*i,M*_. We first consider *r*_*i,M*_, which requires descriptions of the reactions involving the transported species. One such reaction is the synthesis of atRA by choroidal cells, which we assume occurs in the choriocapillaris at a constant rate, *k*_*prod*_ (moles atRA/time/tissue volume), and thus can be described by *k*_*prod*_*a*(*x*). Because of its hydrophobicity, newly-synthesized atRA is transported intracellularly to the cell surface and then is assumed to bind to SA to be transported in the extracellular space. We assume this reaction obeys first order kinetics with equilibrium association constant 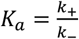

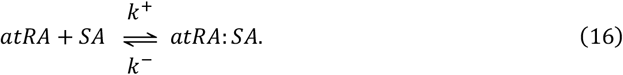

It is of interest to estimate the magnitudes of terms in the above equation. Reported values for *K*_*a*_ range from 3.3 × 10^5^ M^-1^ to 2.3 × 10^6^ M^-1^ [16,31,32] and, as will be seen below, typical SA concentrations in the choroid and sclera are of order 2 × 10^−4^ M. Substitution into equation (16) shows that only about 0.2 to 1.5% of atRA is expected to be unbound at equilibrium (*c*_3_ ≫ *c*_1_). This means that the total amount of atRA (bound and unbound) in a given tissue is well approximated by *c*_3_, making comparison of numerical and experimental results slightly more convenient.

The reaction source term for atRA (species 1) can be written as:

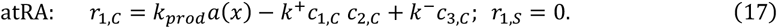

Since atRA cannot be transported unless bound to a carrier, the conservation of mass requires that *r*_1,*C*_ = 0. Equation (17) then immediately implies that

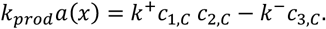

This leads to another simplification: we can replace the term *k*^+^*c*_1,*C*_ *c*_2,*C*_ − *k*^−^*c*_3,*C*_, appearing in the source/sink terms in the transport equations for SA and atRA:SA (species *i* = 2, 3), by *k*_*prod*_*a*(*x*), which we do below.

The degradation of atRA is a complex process that is mediated by CYP26 enzymes [33–37]. To model it we adopt the approach proposed by White et al. [38], who essentially assumed that the cytoplasmic uptake of atRA was proportional to the extracellular concentration of atRA:SA. Following the work of Jing et al. [39], which relates the amount of CYP26 enzymes to the atRA concentration, the concentration of the CYP26 enzymes takes the form 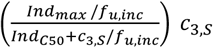, i.e. more atRA leads to greater concentrations of CYP26 (see Supplemental Material §S1.1 for further details). Thus, assuming that the degradation rate of atRA is linearly proportional to the concentration of atRA and the CYP26 enzymes (first order kinetics) [38], we can write the atRA degradation term as

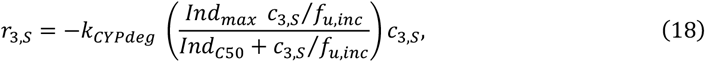

where *k*_*CYPdeg*_ is an effective degradation rate constant that incorporates the effects of all CYP enzymes. The constants *Ind*_*max*_, Ind_*C*50_ and *f*_*u,inc*_ are taken from [39].

In summary, the reaction terms for species 2 and 3 are:

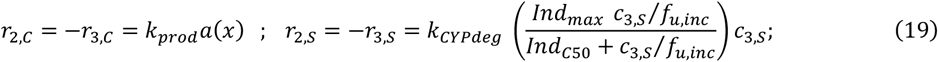

We next consider the transmural transport terms, *b*_*i,M*_. In the sclera there is assumed to be no leakage of species out of vessels, so that *b*_*i*,S_ = 0. However, in the choriocapillaris we must account for transmural transport of SA (species 2) and atRA:SA (species 3), which has been shown to occur by shuttling of caveolae across the capillary endothelial cells [40]. In this case, we assume that the transport can be expressed as a first order rate equation of the form

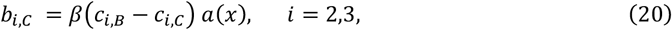

where *β* is a rate constant (units of inverse time) that must be determined by parameter fitting. Similarly to 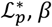, *β* depends on the average surface of the choriocapillaris per unit tissue volume, 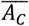. The term *a*(*x*) accounts for the fact that transmural SA and atRA:SA transport occurs only in the choriocapillaris.

##### Boundary conditions for solute transport

We assume that the RPE is “tight” so that there is no permeation of any species (but water) across the RPE, which is written as

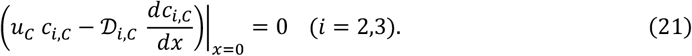

We also require matching of concentrations and mass fluxes at the choroid-sclera interface, i.e.

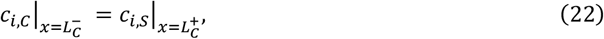

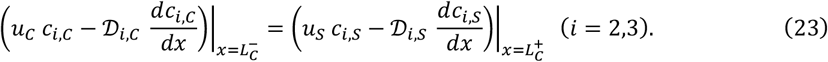

Finally, we assume that there is zero gradient of concentration at the outer scleral surface, i.e.

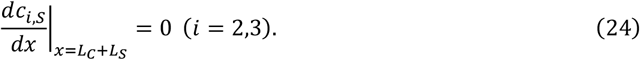

Since we do not reliably know the concentration at the outer scleral surface, we adopt this approach because it provides mathematical closure of the problem while minimally perturbing the concentration field. This condition assumes that solutes in the orbit are well mixed.

A summary of the model and details of the numerical solution approach are described in the Supplemental Material §S1.2 and §S1.3.

### 2.5 Input parameters and sensitivity analysis

The solution of the above system of equations requires specification of input parameter values, some of which are well known and others of which are essentially completely unknown. We grouped these parameters into three categories: those whose values are reasonably well known; those for which some data is available but there is less certainty about their values; and those whose values are extremely uncertain. This grouping is described below; note that all parameter categorizations are for both mouse and human, unless otherwise noted.

- Input parameters that are reasonably well known: *L*_*C*_, *L*_*S*_, *A*_*S*_, *K*_*S*_ in humans, 𝕔, *Q*_*prod*_, *EVP*, 𝒟_*i,C*_ (*i* = 2,3), 𝒟_*i,S*_ (*i* = 2,3), *µ, p*_*orbit*_, c_*i,B*_.
- Input parameters for which some data is available: *K*_*C*_, Δ*P*_*blood*_, Δ*P*_*SCS*_, *α* (required for defining a(x)), u_*RPE*_, *K*_*S*_ in mice, *Ind*_*max*_, Ind_*C*50_, *f*_*u,inc*_.
- Input parameters that are extremely uncertain: *β*, 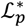, *k*_*prod*_, *k*_*CYPdeg*_.

*Effects of atRA feeding:* atRA feeding in mice leads to a significant increase in plasma atRA:SA concentrations (see Results). We therefore undertook simulations for three sets of conditions:

1. Mice without atRA feeding (control). The atRA:SA concentration in serum, *c*_),−_, was specified based on serum measurements for non-feeding conditions (see below).
2. Mice with atRA feeding. As in case 1, based on measurements from atRA-fed mice.
3. Humans (without atRA feeding). The atRA:SA concentration in the human serum (under no-feeding conditions) was assumed 2.5 nM [41–43].

As previously reported [7], atRA feeding led to a significant increase in atRA concentrations in relevant ocular tissues, with the mean atRA concentration in the RPE+Choroid+Sclera (R/C/S) increasing approximately 100 fold, from 7.5 nM to 0.68 μM.

The model parameters and their values are summarized in Table 1, while we provide a detailed explanation for the choice of these parameter values in the Supplemental Material §S2.3. To find values for parameters which are extremely uncertain, we fit (see Supplemental Material §S2.2) the model outputs to known quantities (IOP, *Q*_*u*_/*Q*_*prod*_, < *c*_*2*_ >_*S*_/*c*_2,*B*_, < *c*_3_ >_*CS*_) reported in the last section of table 1. Here and throughout, the brackets <·> represent a spatial averaging operation, with the subscript following the bracket indicating the domain over which the averaging is occurring, i.e. < *c*_2_ >_*C*_, < *c*_2_ >_*S*_ and < *c*_2_ >_*Cp*_ denote the spatially averaged concentration of SA in the choroid, sclera and choroid + sclera, respectively.

**Table 1.** Model parameters and their values. Details about the source(s) of parameters values can be found in supplemental table S1.

| Parameter Symbol, Description and Units | Values in: |  |
| --- | --- | --- |
|  | Mouse | Human |
| $L_S$ (scleral thickness, $\mu\text{m}$) | 70 | 670 |
| $L_C$ (choroidal thickness, $\mu\text{m}$) | 30 | 268 |
| $\alpha$ (fraction of choroidal thickness occupied by choriocapillaris) | 1/2 | 1/9 |
| $R$ (universal gas constant, J/mol/K) | 8.314 | |
| $T$ (absolute temperature, K) | 310 | |
| $\mu$ (dynamic extravascular viscosity of fluid in choroid/sclera, Pa s) | $0.7 \times 10^{-3}$ | |
| $u_{RPE}$ (pumping velocity of RPE, m/s) | $23 \times 10^{-8}$ | $3 \times 10^{-8}$ |
| $p_{orbit}$ (Pressure in the orbital tissue, mmHg) | 3 | 3 |
| $\Delta P_{blood}$ (Offset between blood pressure in the choroid and IOP, mmHg) | 5 | 5 |
| $\Delta P_{SCS}$ (Offset between IOP and pressure in the suprachoroidal space, mmHg) | 1 | 1 |
| $EVP$ (episcleral venous pressure, mmHg) | 7.5 | 7.1 |
| $Q_{prod}$ (production rate of aqueous humor, $\mu\text{L}/\text{min}$) | $48 \times 10^{-3}$ | 2.53 |
| $\mathbb{C}$ (conventional outflow facility, $\mu\text{L}/\text{min}/\text{mmHg}$) | $5.9 \times 10^{-3}$ | 0.26 |
| $\mathcal{L}_p^*$ (modified hydraulic permeability of choriocapillary walls, 1/s/Pa) | $3.56 \times 10^{-5}$ | $1.84 \times 10^{-7}$ |
| $K_S$ (Darcy permeability of sclera, $\text{m}^2$) | $6.4 \times 10^{-18}$ | $1.33 \times 10^{-18}$ |
| $K_C$ (Darcy permeability of choroid, $\text{m}^2$) | $100 \times K_S$ | $100 \times K_S$ |
| $\mathcal{D}_{2,S}, \mathcal{D}_{3,S}$ (diffusivity of SA and atRA:SA in the sclera, $\text{m}^2/\text{s}$) | $9.6 \times 10^{-12}$ | $9.6 \times 10^{-12}$ |
| $\mathcal{D}_{2,C}, \mathcal{D}_{3,C}$ (diffusivity of SA and atRA:SA in the choroid, $\text{m}^2/\text{s}$) | $8.75 \times 10^{-11}$ | $8.75 \times 10^{-11}$ |
| $\beta$ (rate of SA leakage out of choriocapillary vessels, 1/s) | $1.37 \times 10^{-3}$ | $3.39 \times 10^{-6}$ |
| $c_{2,B} + c_{3,B}$ (total concentration of SA in blood, mM) | 0.41 | 0.68 |
| $c_{3,B}$ (concentration of atRA:SA in blood, mM) | Measured directly (see text) | $2.5 \times 10^{-6}$ |
| $\sigma_i$ (reflection coefficient of solute $i = 2,3$) | 1 | |
| $k_{prod}$ (production [synthesis] rate of atRA by choroidal cells, mM/s) | $7.8 \times 10^{-8}$ | $1.01 \times 10^{-8}$ |
| $k_{CYPdeg}$ (degradation rate of atRA by scleral cells, 1/s) | $3.5 \times 10^{-4}$ | $4.55 \times 10^{-5}$ |
| $Ind_{max}$ (maximum fold induction of atRA degradation) | 33 | |
| $Ind_{c50}$ (atRA concentration at which 50% of the maximum degradation induction occurs, nM) | 90 | |
| $f_{u,inc}$ (fraction unbound in the incubation) | 0.4 | |
| Optimization targets |  |  |
| $IOP$ (intraocular pressure, mmHg) | 14.4 | 14.9 |
| $Q_u/Q_{prod}$ (% of outflow that is unconventional) | 20% | 20% |
| $\langle c_2 \rangle_S / c_{2,B}$ (average SA concentration in the sclera normalized by serum SA concentration) | 19% | 19% |
| $\langle c_3 \rangle_{CS}$ (mean concentration of atRA:SA in the choroid and sclera, mM) | Control: $0.75 \times 10^{-5}$<br>Feeding: $0.68 \times 10^{-3}$ | N/A |

In order to investigate the effect of the uncertainty of the model parameters we performed global sensitivity analysis using the Fourier amplitude sensitivity test (eFAST) [44]. In this method, all considered parameters are varied over their corresponding ranges (see Supplemental Material §S3). A total sensitivity index is then generated for each parameter, which is a measure of the importance of that parameter in determining the value of the observed model output and which also accounts for interactions of the parameter with other parameters. The higher the value of the sensitivity index, the greater the parameter’s impact on the model output. In the sensitivity analysis we considered the following parameters:

- *K*_*S*_, *u*_*RPE*_, *Ind*_*C*50_ and *c*_2,*B*_ varying over the range [0.5,1.5] times their reference values in table 1;
- Δ*P*_*blood*_ ∈ [3,7] mmHg and Δ*P*_*SCS*_ ∈ [0,2] mmHg; and
- The extremely uncertain parameters (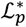, *β, k*_*prod*_, *k*_*CYcdeg*_) varying over the range of [1/3,3] times their reference values.

## 3. Results

We first present model predictions regarding only fluid and SA transport, i.e. ignoring atRA, since its concentrations are negligible and thus do not affect the transport of other species. This helps us understand this complex system, necessary to investigate the dynamics of atRA transport and its dependence on fluid flow and SA transport. We then examine the combined fluid, SA, and atRA transport, starting with experimental measurements of atRA serum concentrations, followed by the results of the combined transport.

### 3.1 Fluid and SA transport

Simulations using parameter values in table 1 aligned well with available experimental measurements (see table 2), even if we recognize that for some of these measurements there is significant uncertainty. Using optimized values for *β* and 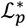, we obtained an *IOP* of approximately 14.9 mmHg in humans, with slightly less than 20% of outflow leaving the eye via the unconventional route; both predictions are close to the target values used in the optimization. The scleral SA concentration was about 18% of the blood concentration, close to the target of 19%. The model further predicted that ~55.5d of the fluid drains via choroidal blood vessels (uveovortex flow), while the remaining ~11.5d drains through the sclera (uveoscleral flow).

**Table 2.** Outputs from the input parameter optimization step for humans and mice without atRA feeding. Values in parentheses in the first three rows represent target values during the optimization of input parameter values, as listed in Table 1. Quantities in the last three rows were not optimization targets but were computed due to their physiologic interest. See text for descriptions and definitions of individual parameters. Note that 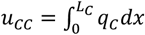.

| Parameter | Mouse | Human |
| --- | --- | --- |
| *IOP* (mmHg) | 14 (14.4) | 14.9 (14.9) |
| Unconventional outflow as a percentage of total outflow, $Q_u/Q_{prod}$ (%) | 19.9 (20) | 19.9 (20) |
| Average SA concentration in the sclera as a fraction of SA concentration in blood, $\langle c_2 \rangle_s / c_{2,B}$ | 0.19 (0.19) | 0.18 (0.19) |
| Fluid outflow rate across the sclera per unit area, $u_s$ (m/s) | $1.75 \times 10^{-7}$ | $0.41 \times 10^{-8}$ |
| Fluid flow rate out of the choroidal capillaries per unit area, $u_{cc}$ (m/s) | $-0.6 \times 10^{-7}$ | $-3.16 \times 10^{-8}$ |
| Serum albumin leakage rate from the choroidal capillaries, mol/s | $4.34 \times 10^{-13}$ | $7.46 \times 10^{-13}$ |

Further insight into the model’s behavior is gained by graphing predicted pressure and serum albumin concentration in the choroid and sclera for the human case (figure 2a). Within the choroid, the pressure is essentially constant, due to the high choroidal hydraulic permeability compared to the sclera. This also implies that, based on equation (7), the pressure in the choroidal tissue is everywhere approximately equal to *IOP* − Δ*P*_*SCS*_. As predicted by equation (4), in the sclera, where there are no exchanges of fluid between tissue and blood vessels, the pressure decreases linearly from its value in the choroid to the orbital pressure (3 mmHg).

**Figure 2:**
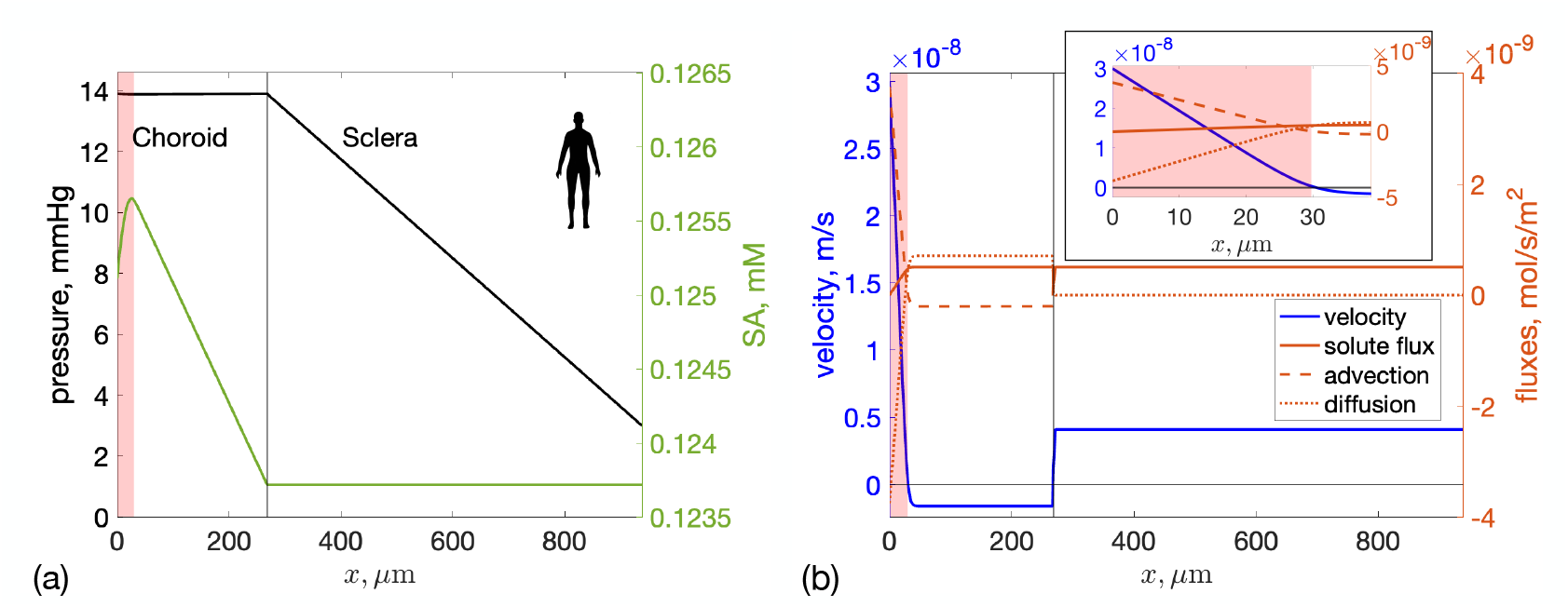
Model predictions of fluid and serum albumin (SA) transport in the human eye. The left edge of the plot (*x* = 0) represents the position of the RPE, the black vertical line indicates the boundary between the choroid and sclera (*x* = *L*_*C*_) and the right edge is the external surface of the sclera (*x* = *L*_*C*_ + *L*_*S*_). The choriocapillaris is indicated by red shading. **(a)** Left axis: Pressure distribution (black), showing almost no pressure drop across the choroid and a constant pressure gradient across the sclera. Right axis: SA concentration distribution (green). **(b)** Left axis: Fluid flux in the choroid and sclera. Horizontal black line represents zero velocity. The small decrease in fluid velocity just to the right of the choriocapillaris is due to the assumed spatial distribution of filtering capillaries being a smoothed step function (equation (8)), which causes some fluid exchange between tissue and blood vessels in a very thin zone outside of the shaded region. Right axis: Total, advective and diffusive fluxes of SA (red). Velocities and fluxes are taken as positive in the external direction, i.e. from the RPE towards the orbit. Inset: a zoomed-in view focusing on the choriocapillaris.

Figure 2a also shows the spatial distribution of SA in the choroid and sclera. The SA concentration increases in the choriocapillaris as SA leaks from vessels into the tissue, then decreases linearly in the outer choroid and remains constant in the sclera. Total SA flux is shown in figure 2b, with advective and diffusive components shown separately. The transport in the choroid is diffusion dominated, while the transport in the sclera is purely due to advection. This explains the uniform profile of SA in the sclera in figure 2a.

Figure 2b also shows the predicted fluid velocity in the choroid and sclera. At the RPE (*x* = 0) the velocity takes the imposed value due to RPE pumping and then rapidly decreases, as fluid is adsorbed into the choriocapillaris. At the choroid/sclera interface, i.e. in the SCS, the model predicts a jump in fluid velocity, which is due to the unconventional flow in the SCS. In both the sclera and the region of the choroid outside the choriocapillaris, the velocity is constant, due to mass conservation (equation (3)). The velocity in the sclera is positive, indicating that fluid is exiting the eye through the sclera. The velocity in the external region of the choroid is slightly negative, which implies that part of the unconventional flow is drained into the choroidal blood vessels (uveovortex flow), although the majority crosses the sclera (uveoscleral flow).

The last column of table 2 and figure 3 report results for the mouse eye, which are qualitatively similar to the human. It is worth noticing, however, that the model predicts a positive fluid velocity everywhere in the choroid and sclera, which implies that the choroidal capillaries do not drain all the fluid that enters the choroid through the RPE and that the unconventional aqueous outflow exits the eye entirely through the sclera. In fact, the ratio of fluid (from both the RPE and the SCS) drained through the choriocapillaris to that through the sclera is 0.34 in mice, compared to 7.7 in humans, due to the mouse sclera being thinner and more permeable to water.

**Figure 3:**
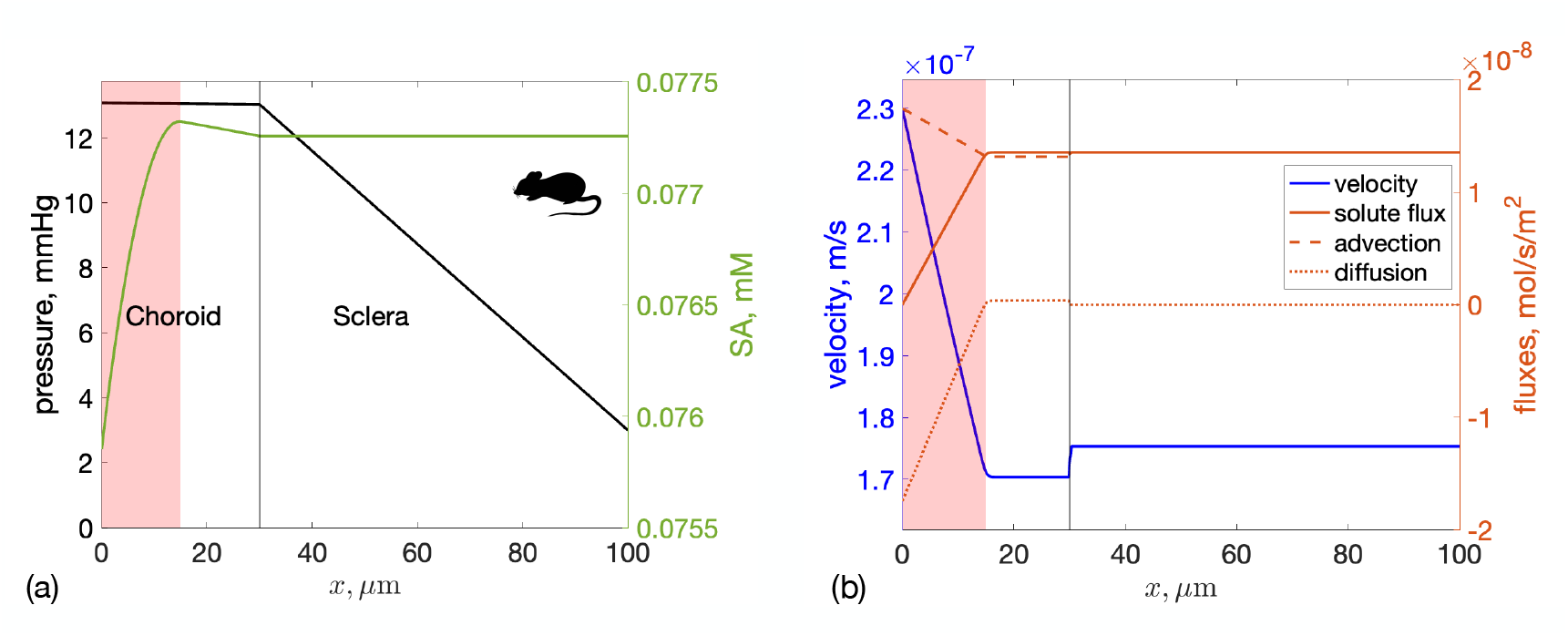
Fluid and albumin transport in the mouse eye. The plotted quantities are the same as for figure 2, although please note that the spatial scale (*x*-axis) and vertical scales (*y*-axes) are different than in the human eye.

We finally note that the rate of SA leakage from blood vessels (last row in table 2) in both mouse and human have similar magnitudes to the value of 9.6 × 10^−13^ mol/s measured by Bill [45] in rabbits, which is reassuring.

### 3.2 Sensitivity analysis for fluid and SA transport

A sensitivity analysis identifies which input parameters most influence model outputs. We conducted a global sensitivity analysis on fluid and SA transport for both humans and mice, as shown in figures 4a and 4b. Interestingly, the sensitivity of each input parameter differed appreciably between the human and the mouse.

**Figure 4:**
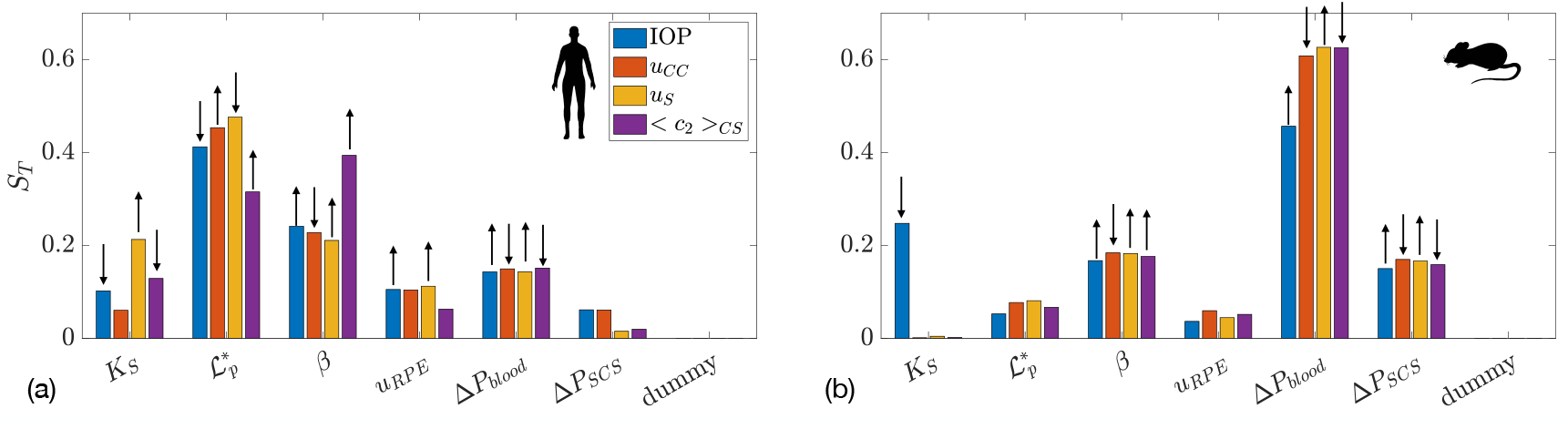
Total sensitivity index for four key outcome measures: IOP; fluid outflow rate into the choriocapillaries (*u*_*CC*_) and the sclera (*u*_*S*_); and average albumin concentration in the choroid and sclera (< *c*_*2*_ >_*CS*_) in: **(a)** human and **(b)** mouse. Bars indicate the sensitivity of each input parameter to a specific model output. The arrow above each bar indicates how the parameter affects the model output: an upward pointing arrow indicates that an increase in the parameter value would, on average, induce an increase in the model output, and *mutatis mutandis*. The input parameters are: *K*_*S*_ – Darcy permeability of the sclera, 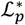-modified hydraulic permeability of the choriocapillary walls, *β* - rate of SA leakage out of choriocapillary vessels, *u*_*RPE*_ – RPE pumping velocity, Δ*P*_*blood*_ - offset between blood pressure in the choroid and IOP, and Δ*P*_*SCS*_ – offset between IOP and pressure in the suprachoroidal space.

Let us consider the human case first, where figure 4a shows that the hydraulic permeability of the walls of the choroidal capillaries, 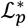, had a strong effect on all the considered model outputs: as 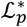 went up, fluid flux into the choroidal capillaries increased because the resistance of the uveovortex pathway decreased. As a consequence, both IOP and the uveoscleral portion of unconventional outflow also decreased. The permeability of the choriocapillaris to SA, *β*, was the most relevant determinant of SA concentration, since SA enters the domain only through blood vessels. The scleral hydraulic permeability, *K*_*S*_, significantly affected the amount of fluid flow through the sclera. Among the other parameters considered, the least relevant for all model outputs were the RPE pumping flux, *u*_*RPE*_, and the difference between IOP and the suprachoroidal space pressure, Δ*P*_*SCS*_.

In the mouse (figure 4b) the parameter that most affected all model outputs was the difference between blood and choroidal pressures, Δ*P*_*blood*_. Interestingly, scleral permeability, *K*_*S*_, only affected *IOP*, not other outputs. This is likely because, within the parameter range studied, all water flux occurred through the uveoscleral pathway, which had much lower resistance than the uveovortex pathway, keeping scleral drainage constant. This was different than the situation in human, where the choroid was much more effective at fluid drainage than in the mouse so that changes in choroidal parameters resulted in an appreciable change in model outputs.

### 3.3 Measurements of atRA concentration in serum

The concentration of atRA in serum for mice after feeding is shown in Figure 5. We measured a rapid and significant increase in serum levels of atRA, from 1.25 ± 0.54 pmol/mL (mean ± standard deviation, all time points pooled, n=8 samples) in control animals to 4.1 nmol/mL (n=2 samples) at 30 minutes after feeding. Serum atRA concentrations continued to increase with time, reaching a plateau of approximately 12 nmol/mL that persisted out to 120 minutes after feeding. We note that the increase in plasma atRA concentration 30 minutes after feeding followed by a plateau is similar to that seen in mice receiving oral 4-oxo-trans-RA, an oxidative metabolite of retinoic acid [46].

**Figure 5:**
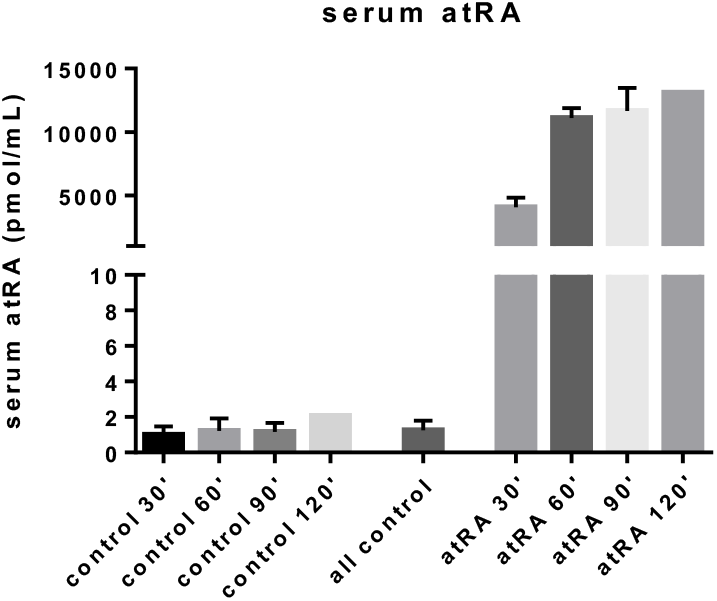
Measured serum concentrations of atRA in control mice and in atRA-fed mice at various time points after completion of atRA feeding. The “all control” bar represents the average over all control cohorts. Plotted quantities are mean ± standard deviation. Note that the control and atRA 120 minute cohorts consisted of only a single mouse each, precluding computation of a standard deviation.

Based on these measurements, in the model we imposed the concentration of atRA:SA in serum, *c*_),−_, to be 1.25 pmol/mL and 11.67 nmol/mL (mean atRA concentration 90 minutes after feeding) for non-feeding and feeding conditions, respectively. We also investigate the effect of this parameter in the sensitivity analysis.

### 3.4 Predicted atRA concentrations in selected ocular tissues

atRA transport within the choroid and sclera is complex, depending on a combination of advection due to the fluid flow (as described above), atRA diffusion, atRA synthesis in the choroid, and atRA degradation in the sclera. We explore this by plotting SA and atRA:SA concentration profiles (figures 6a-c), and atRA consumption by the sclera (figure 6d). Relevant integrated quantities, such as atRA synthesis rate, vessel leakage rate, degradation rate, and mean scleral atRA:SA concentration, are presented in table 3.

**Table 3.**
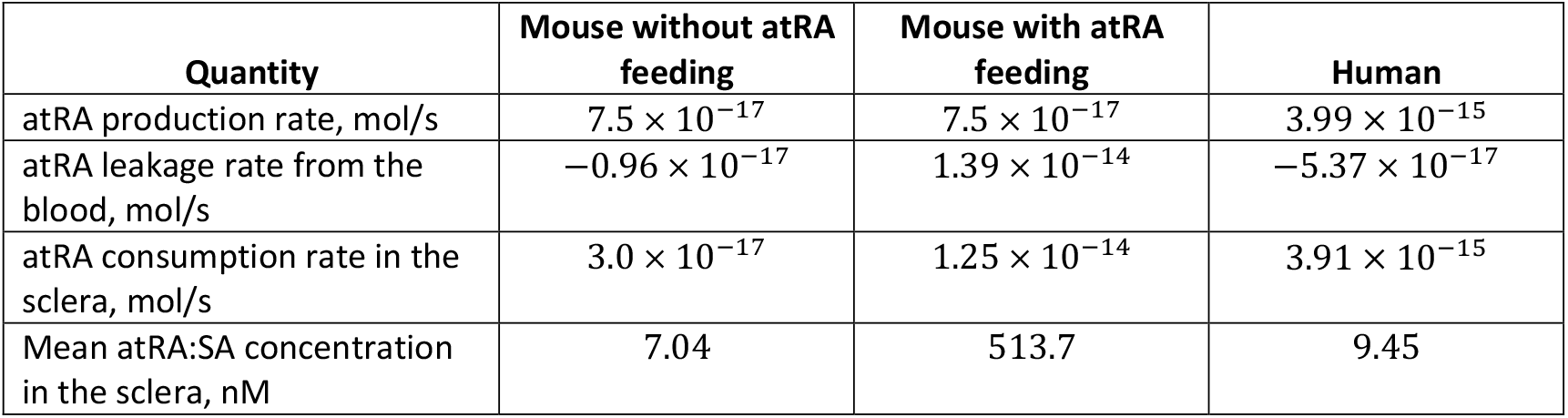
atRA production and consumption rates and mean atRA:SA concentration in the sclera, as computed by the model in mouse (with and without atRA feeding) and in human. The quantities in the rows are: total atRA cellular synthesis rate, 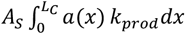 total atRA inflow (leakage) rate from the blood to the extravascular space, 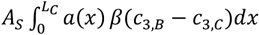 total atRA consumption rate in the entire sclera, 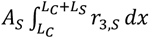 and average concentration of atRA in the sclera < *c*_3_ >_*s*_.

**Figure 6:**
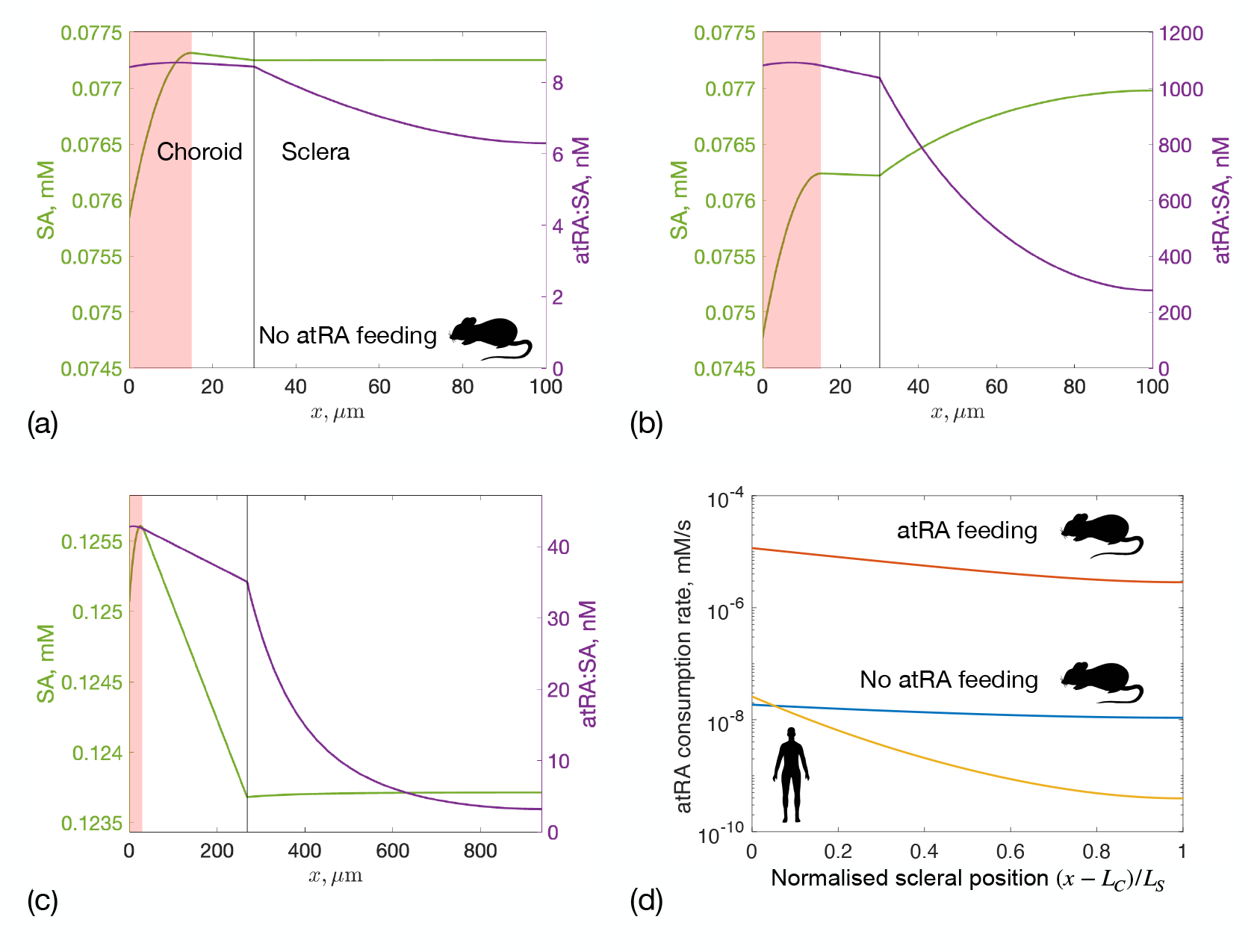
SA (left axis) and atRA:SA (right axis) concentrations in the choroid and sclera for: **(a)** a mouse without atRA feeding, **(b)** a mouse with atRA feeding, and **(c)** a human. Note the different axis scales in panels (a)-(c). The sum of the two curves in each of panels (a) through (c) gives the curves reported in Figure 3(a), i.e. the total albumin concentration. Panel **(d)** shows the atRA consumption rate per unit volume of tissue within the sclera, *r*_3,*S*_. The *x*-axis in panel (d) is the normalized distance from the choroid-sclera interface to the exterior sclera, i.e. choroid is on the left and the orbit is on the right. This approach allows us to create a single graph that includes data from both mouse and human. The three curves in panel (d) correspond to: mouse without atRA feeding (blue, max(*r*_3,*S*_) = 1.85 × 10^−8^ mM/s), mouse with atRA feeding (red, max(*r*_3,*S*_) = 1.16 × 10^−5^ mM/s) and human (yellow, max(*r*_3,*S*_) = 2.6 × 10^−8^ mM/s).

In mice, atRA:SA concentrations in the choroid and sclera are much higher under feeding conditions (by approximately 2 orders of magnitude) compared to non-feeding conditions (figure 6a vs. 6b). Under feeding conditions, atRA:SA leakage from the bloodstream is ~186 times larger than atRA synthesis by choroidal cells (table 3). In all cases, the atRA:SA concentration increases across the choroicapillaris (shaded area in figures 6a and 6b), where atRA is either locally synthesized or leaks from the vessels, and then remains approximately constant in the outer choroid. In the sclera, atRA:SA concentration decreases due to degradation by scleral fibroblasts. However, in the mouse under non-feeding conditions, there is an imbalance between atRA production and consumption in the sclera (only 40% of atRA is consumed) resulting in small spatial variability in atRA:SA concentration (and also atRA consumption). This means that a substantial amount of atRA exits into the orbit. Interestingly, in the non-feeding case, a small amount of atRA:SA is predicted to leak back into the choroidal circulation, since the concentration of atRA:SA is higher in the insterstitium than in the vessels due to atRA production in the choriocapillaris.

Despite the decrease in atRA:SA concentration across the sclera, scleral concentrations remain in the 3-40 nM range for both non-feeding mice and humans. For example in humans, the average scleral atRA:SA concertation is 9.45 nM vs. 7.04 nM in the mouse without atRA feeding (table 3). Interestingly, at the sclera’s external boundary, atRA:SA concentrations are similar in humans (3.2 nM) and mice (6.3 nM) under non-feeding conditions, even though 98% of synthesized atRA is degraded in the human sclera. Under feeding conditions in the mouse, atRA levels in the sclera jump by nearly 2 orders of magnitude, and a significant concentration of atRA:SA (278 nM) is predicted to be carried into the orbit by fluid crossing the sclera, which may be experimentally testable. Finally, under feeding conditions, high atRA levels impact free SA profiles, as atRA:SA concentrations become comparable to free SA (figure 6b). This means that, as atRA degrades, atRA:SA decreases and free SA increases.

It is of interest to compute spatial profiles of atRA consumption rates across the sclera (figure 6d), since this quantity represents atRA that has entered cells where it can exert biological effect. In the atRA-fed mouse, the consumption profile decreases much faster than in the non-feeding case, since more atRA is consumed by the sclera. In the human (yellow), the consumption rate decreases by almost two orders of magnitude from the inner to the outer sclera.

### 3.5 Sensitivity analysis for atRA transport

In figure 7 we show the results of global sensitivity analysis when atRA transport is considered. Figures 7(a,c) show the total sensitivity index for mean atRA:SA concentration in the sclera and in the orbit, in the non-atRA-fed mouse and human, respectively. Unsurprisingly, the most important parameters are the production of atRA, *k*_*prod*_, and its degradation rate constant, *k*_*CYPdeg*_. In the mouse, the parameter that influences the exchange with choriocapillaris, *β*, is also important: increasing *β* increases the outflow of atRA:SA into the blood, thus reducing atRA:SA in the sclera. This is not the case for the human, however, as the outflow into the capillaries is two orders of magnitude smaller than atRA production (table 2). To gain additional insight, in figure S1 we show atRA levels in the sclera as a function of atRA degradation and production rates.

**Figure 7:**
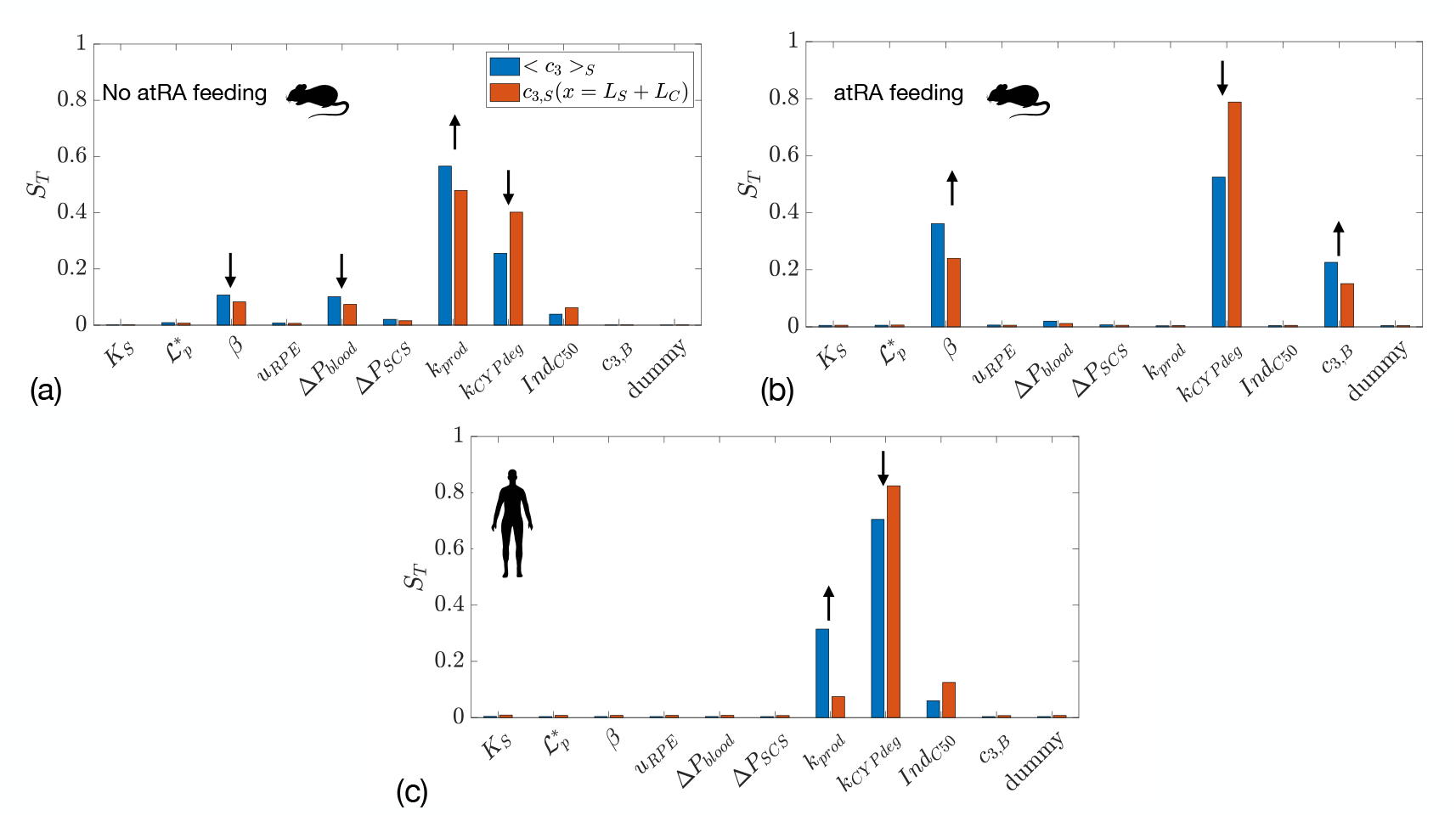
Total sensitivity index (*S*_*T*_) for two outcome measures: mean atRA:SA concentration in the sclera < *c*_3_ >_*s*_ and atRA:SA concentration in the orbit *c*_3,*S*_(*x* = *L*_*S*_ + *L*_*C*_) in: (a) mice without atRA feeding; (b) mice with atRA feeding; and (c) humans. The first 6 quantities on the *x*-axis are the same as in Figure 4, with the addition of *k*_*prod*_ (atRA production rate by choroidal cells), *k*_*CYPdeg*_(the atRA degradation rate constant), *Ind*_*C*50_ (the atRA concentration at which 50% of the maximum atRA degradation induction occurs), and *c*_3,*B*_ (the concentration of atRA:SA in plasma).

The case of atRA feeding in the mouse is shown in figure 7(b), where the atRA production rate is no longer important, since local production is dwarfed by atRA leakage from the vasculature. Thus, the key parameters here are atRA:SA concentration in blood, *c*_3,*B*_, the rate of (atRA:)SA leakage out of choriocapillary vessels, *β*, and the degradation rate constant, *k*_*CYPdeg*_.

## 4. Discussion

The major finding of this work is that, in both mouse and human eyes, atRA produced by stromal cells in the inner choroid is predicted to permeate into and across the sclera at concentrations that are biologically active (see below). Further, atRA bound to serum albumin (atRA:SA) in choroidal blood is predicted to be able to extravasate from the choriocapillaris and be carried into and across the sclera at elevated concentrations, relevant to feeding experiments in mice. These observations are consistent with atRA having an important role in retinoscleral signaling during myopigenesis.

More specifically, in both human and mouse eyes under non-feeding conditions, we predict scleral atRA concentrations in the ~3-40 nM range. The central question is then: are these atRA levels biologically significant? This is not a trivial question to answer, but some insight can be gained by noting that the EC50 concentration for atRA binding to RARβ is 9 nM, as determined by a luciferase reporter assay in cell culture [47]. We selected RARβ because this is reported to be the predominant retinoic acid receptor isoform present in scleral fibroblasts. Evidently more study is required, but it does appear that atRA predicted concentrations in the sclera lie in a biologically significant range.

To make our predictions we have employed a combination of mathematical modeling and direct measurement of (bulk tissue) atRA concentrations. Unfortunately, we cannot currently make quantitative, spatially-resolved atRA concentration measurements within the relevant mouse ocular tissues. However, mathematical modeling helps us to assess spatial variability in atRA concentrations, to understand the relative importance of different atRA transport mechanisms, and to study the effects of parameters on atRA transport. Evidently, such modeling requires a large number of input parameters, some of which are incompletely or entirely unknown. We have therefore resorted to fitting input parameter values, and to employing a sensitivity analysis to identify those input parameters that are most influential in our model. Future work should consider making direct, quantitative measurements of some of these most influential parameters, although that will likely be technically challenging.

It is clear that atRA transport in the eye is complex, involving endogenous synthesis, leakage from vessels, diffusion, advection and degradation. Importantly, the model predicts that atRA transport depends critically on fluid and SA transport, which essentially required us to develop a novel model of unconventional flow, even though this was not the original goal. The comparison between mice and humans has revealed a number of interesting features. Specifically, there are fundamental differences in fluid and SA transport between mouse and human, largely driven by the much thinner mouse sclera which causes the uveoscleral component of unconventional outflow to be much more significant than the uveovortex component in the mouse eye (vs. in the human eye). This suggests that the utility of the mouse as a model for human unconventional outflow may have previously unappreciated limitations.

One interesting prediction of our model is that, under experimental atRA feeding conditions in mice, a significant amount of atRA enters the orbit by transport across the sclera. This is in principle an experimentally testable prediction of the model, although feeding will of course elevate atRA levels in the vasculature of the orbit which could confound atRA measurements in orbital tissues.

This work is subject to certain limitations, among which we note the following:

- The model is simplified, inasmuch as it does not include all possible signaling pathways, considers only one-dimensional transport, and simplifies some complex biological processes, e.g. atRA degradation. However, we feel strongly that having a more complex model was unwarranted at the present time due to the limited amount of experimental data that was available. Indeed, some of the available experimental data shows significant scatter, and some data is only available in certain species, e.g. in mouse only or rabbit only. Our sensitivity analysis, showing which parameters are important, is particularly relevant in this context since it will hopefully motivate targeted experimental measurements. As better experimental data becomes available on retinoic acid signaling and degradation, we look forward to refining our model.
- We were forced to use scaling arguments to translate production and degradation rates of atRA and SA transport rates from mice to human. Future work should study the applicability of such scaling for atRA synthesis and degradation.
- In this model we do not account for any possible signaling that lies interior to the RPE. Such signaling is clearly an important aspect of the retinoscleral signaling cascade and is a subject of ongoing research.
- In our model we do not explicitly account for intracellular transport or binding, e.g. as mediated by CRABP2 [48,49].
- atRA feeding in mice yields very high levels of atRA, which might overwhelm some regulatory and transport mechanisms, which is not included to the model.
- The model is steady and therefore does not consider transient effects occurring during myopigenesis. This assumption does not necessarily hold if animals are fed atRA: there is a time-dependent change in atRA serum levels, which reach a peak after feeding and persist at that level for at least several hours. Additionally, there is temporal variability in atRA levels during myopia induction and recovery, which is not included in the model. Future work should include a dynamic component that accounts for these dynamic processes.

## Supporting information

Experimental data

## Acknowledgements

We gratefully acknowledge financial support from NIH F32 EY035573 (MBF), NIH T32 EY007092 (MBF), NIH R01 EY016435 (MTP), NIH R01 EY033361 (MTP, CRE, MAK) and NIH P30 EY006360; Dept. of Veterans Affairs Research Career Scientist Award RX003134 (MTP); RPB Challenge Grant to Emory Ophthalmology; Georgia Research Alliance (CRE); and University of Maryland School of Pharmacy Mass Spectrometry Center (SOP1841-IQB2014). Prof. Ethier participated in the Visiting Researcher and Visiting Professor Funding Schemes from the University of Genoa, in the years 2022, 2023 and 2024.

## References

1. Holden BA, Fricke TR, Wilson DA, Jong M, Naidoo KS, Sankaridurg P, Wong TY, Naduvilath TJ, Resnikoff S. 2016 Global Prevalence of Myopia and High Myopia and Temporal Trends from 2000 through 2050. Ophthalmology 123, 1036–1042. (doi:10.1016/j.ophtha.2016.01.006)

2. Baird PN et al. 2020 Myopia. Nat. Rev. Dis. Primer 6, 99. (doi:10.1038/s41572-020-00231-4)

3. Buch H, Vinding T, Nielsen NV. 2001 Prevalence and causes of visual impairment according to World Health Organization and United States criteria in an aged, urban Scandinavian population: the Copenhagen City Eye Study. Ophthalmology 108, 2347–2357. (doi:10.1016/s0161-6420(01)00823-5)

4. Jonas JB, Jonas RA, Bikbov MM, Wang YX, Panda-Jonas S. 2023 Myopia: Histology, clinical features, and potential implications for the etiology of axial elongation. Prog. Retin. Eye Res. 96, 101156. (doi:10.1016/j.preteyeres.2022.101156)

5. Brown DM, Mazade R, Clarkson-Townsend D, Hogan K, Datta Roy PM, Pardue MT. 2022 Candidate pathways for retina to scleral signaling in refractive eye growth. Exp. Eye Res. 219, 109071. (doi:10.1016/j.exer.2022.109071)

6. Gentle A, McBrien NA. 2002 Retinoscleral control of scleral remodelling in refractive development: a role for endogenous FGF-2? Cytokine 18, 344–348. (doi:10.1006/cyto.2002.1046)

7. Brown DM, Yu J, Kumar P, Paulus QM, Kowalski MA, Patel JM, Kane MA, Ethier CR, Pardue MT. 2023 Exogenous All-Trans Retinoic Acid Induces Myopia and Alters Scleral Biomechanics in Mice. Invest. Ophthalmol. Vis. Sci. 64, 22. (doi:10.1167/iovs.64.5.22)

8. Mao J-F, Liu S-Z, Dou X-Q. 2012 Retinoic acid metabolic change in retina and choroid of the guinea pig with lens-induced myopia. Int. J. Ophthalmol. 5, 670–674. (doi:10.3980/j.issn.2222-3959.2012.06.04)

9. Summers JA. 2019 Retinoic Acid in Ocular Growth Regulation. In Vitamin A (eds L Queiroz Zepka, V Vera de Rosso, E Jacob-Lopes), IntechOpen. (doi:10.5772/intechopen.84586)

10. Summers JA, Cano EM, Kaser-Eichberger A, Schroedl F. 2020 Retinoic acid synthesis by a population of choroidal stromal cells. Exp. Eye Res. 201, 108252. (doi:10.1016/j.exer.2020.108252)

11. Mertz JR, Wallman J. 2000 Choroidal Retinoic Acid Synthesis: A Possible Mediator between Refractive Error and Compensatory Eye Growth. Exp. Eye Res. 70, 519–527. (doi:10.1006/exer.1999.0813)

12. Morgan IG. 2003 The biological basis of myopic refractive error. Clin Exp Optom 86, 276–288.

13. Niederreither K, Dollé P. 2008 Retinoic acid in development: towards an integrated view. Nat. Rev. Genet. 9, 541–553. (doi:10.1038/nrg2340)

14. Ghyselinck NB, Duester G. 2019 Retinoic acid signaling pathways. Dev. Camb. Engl. 146, dev167502. (doi:10.1242/dev.167502)

15. Liang C et al. 2021 Overview of all-trans-retinoic acid (ATRA) and its analogues: Structures, activities, and mechanisms in acute promyelocytic leukaemia. Eur. J. Med. Chem. 220, 113451. (doi:10.1016/j.ejmech.2021.113451)

16. N’soukpoé-Kossi CN, Sedaghat-Herati R, Ragi C, Hotchandani S, Tajmir-Riahi HA. 2007 Retinol and retinoic acid bind human serum albumin: stability and structural features. Int. J. Biol. Macromol. 40, 484–490. (doi:10.1016/j.ijbiomac.2006.11.005)

17. Lixa C, Clarkson MW, Iqbal A, Moon TM, Almeida FCL, Peti W, Pinheiro AS. 2019 Retinoic Acid Binding Leads to CRABP2 Rigidification and Dimerization. Biochemistry 58, 4183–4194. (doi:10.1021/acs.biochem.9b00672)

18. Norris AW, Cheng L, Giguère V, Rosenberger M, Li E. 1994 Measurement of subnanomolar retinoic acid binding affinities for cellular retinoic acid binding proteins by fluorometric titration. Biochim. Biophys. Acta 1209, 10–18. (doi:10.1016/0167-4838(94)90130-9)

19. Gottesman ME, Quadro L, Blaner WS. 2001 Studies of vitamin A metabolism in mouse model systems. BioEssays 23, 409–419. (doi:10.1002/bies.1059)

20. Summers JA, Harper AR, Feasley CL, Van-Der-Wel H, Byrum JN, Hermann M, West CM. 2016 Identification of Apolipoprotein A-I as a Retinoic Acid-binding Protein in the Eye. J. Biol. Chem. 291, 18991–19005. (doi:10.1074/jbc.M116.725523)

21. Chatterjee S, Roy A, Jianshi Y, Read AT, Bentley-Ford MR, Pardue MT, Kane MA, Finn MG, Ethier CR. In press. Binding partners for all-trans retinoic acid and implications for myopigenesis. BioRXiv (doi:10.1101/2025.01.04.631331)

22. Mertz JR, Wallman J. 2000 Choroidal Retinoic Acid Synthesis: A Possible Mediator between Refractive Error and Compensatory Eye Growth. Exp. Eye Res. 70, 519–527. (doi:10.1006/exer.1999.0813)

23. Harper AR, Wang X, Moiseyev G, Ma J-X, Summers JA. 2016 Postnatal Chick Choroids Exhibit Increased Retinaldehyde Dehydrogenase Activity During Recovery From Form Deprivation Induced Myopia. Investig. Opthalmology Vis. Sci. 57, 4886. (doi:10.1167/iovs.16-19395)

24. Johnson M, McLaren JW, Overby DR. 2017 Unconventional aqueous humor outflow: A review. Exp. Eye Res. 158, 94–111. (doi:10.1016/j.exer.2016.01.017)

25. Brubaker RF. 2004 Goldmann’s equation and clinical measures of aqueous dynamics. Exp. Eye Res. 78, 633–637. (doi:10.1016/j.exer.2003.07.002)

26. Jones JW, Pierzchalski K, Yu J, Kane MA. 2015 Use of fast HPLC multiple reaction monitoring cubed for endogenous retinoic acid quantification in complex matrices. Anal. Chem. 87, 3222–3230. (doi:10.1021/ac504597q)

27. Brown DM, Kowalski MA, Paulus QM, Yu J, Kumar P, Kane MA, Patel JM, Ethier CR, Pardue MT. 2022 Altered Structure and Function of Murine Sclera in Form-Deprivation Myopia. Invest. Ophthalmol. Vis. Sci. 63, 13. (doi:10.1167/iovs.63.13.13)

28. Kane MA, Napoli JL. 2010 Quantification of Endogenous Retinoids. Methods Mol. Biol. Clifton NJ 652, 1–54. (doi:10.1007/978-1-60327-325-1_1)

29. Emi K, Pederson JE, Toris CB. 1989 Hydrostatic pressure of the suprachoroidal space. Invest. Ophthalmol. Vis. Sci. 30, 233–238.

30. M D Aje F, Ea GR R. 2020 Fluid and solute transport across the retinal pigment epithelium: a theoretical model. J. R. Soc. Interface 17. (doi:10.1098/rsif.2019.0735)

31. Maiti TK, Ghosh KS, Debnath J, Dasgupta S. 2006 Binding of all-trans retinoic acid to human serum albumin: Fluorescence, FT-IR and circular dichroism studies. Int. J. Biol. Macromol. 38, 197–202. (doi:10.1016/j.ijbiomac.2006.02.015)

32. Belatik A, Hotchandani S, Bariyanga J, Tajmir-Riahi HA. 2012 Binding sites of retinol and retinoic acid with serum albumins. Eur. J. Med. Chem. 48, 114–123. (doi:10.1016/j.ejmech.2011.12.002)

33. Topletz AR, Tripathy S, Foti RS, Shimshoni JA, Nelson WL, Isoherranen N. 2015 Induction of CYP26A1 by Metabolites of Retinoic Acid: Evidence That CYP26A1 Is an Important Enzyme in the Elimination of Active Retinoids. Mol. Pharmacol. 87, 430–441. (doi:10.1124/mol.114.096784)

34. Isoherranen N, Zhong G. 2019 Biochemical and physiological importance of the CYP26 retinoic acid hydroxylases. Pharmacol. Ther. 204, 107400. (doi:10.1016/j.pharmthera.2019.107400)

35. Tay S, Dickmann L, Dixit V, Isoherranen N. 2010 A Comparison of the Roles of Peroxisome Proliferator-Activated Receptor and Retinoic Acid Receptor on CYP26 Regulation. Mol. Pharmacol. 77, 218–227. (doi:10.1124/mol.109.059071)

36. Topletz AR, Thatcher JE, Zelter A, Lutz JD, Tay S, Nelson WL, Isoherranen N. 2012 Comparison of the function and expression of CYP26A1 and CYP26B1, the two retinoic acid hydroxylases. Biochem. Pharmacol. 83, 149–163. (doi:10.1016/j.bcp.2011.10.007)

37. Ghaffari H, Petzold LR. 2018 Identification of influential proteins in the classical retinoic acid signaling pathway. Theor. Biol. Med. Model. 15, 16. (doi:10.1186/s12976-018-0088-7)

38. White RJ, Nie Q, Lander AD, Schilling TF. 2007 Complex regulation of cyp26a1 creates a robust retinoic acid gradient in the zebrafish embryo. PLoS Biol. 5, e304. (doi:10.1371/journal.pbio.0050304)

39. Jing J, Nelson C, Paik J, Shirasaka Y, Amory JK, Isoherranen N. 2017 Physiologically Based Pharmacokinetic Model of All-trans-Retinoic Acid with Application to Cancer Populations and Drug Interactions. J. Pharmacol. Exp. Ther. 361, 246–258. (doi:10.1124/jpet.117.240523)

40. Nakanishi M, Grebe R, Bhutto IA, Edwards M, McLeod DS, Lutty GA. 2016 Albumen Transport to Bruch’s Membrane and RPE by Choriocapillaris Caveolae. Invest. Ophthalmol. Vis. Sci. 57, 2213–2224. (doi:10.1167/iovs.15-17934)

41. Jones JW, Pierzchalski K, Yu J, Kane MA. 2015 Use of fast HPLC multiple reaction monitoring cubed for endogenous retinoic acid quantification in complex matrices. Anal. Chem. 87, 3222–3230. (doi:10.1021/ac504597q)

42. Wyss R, Bucheli F. 1997 Determination of endogenous levels of 13-cis-retinoic acid (isotretinoin), all-trans-retinoic acid (tretinoin) and their 4-oxo metabolites in human and animal plasma by high-performance liquid chromatography with automated column switching and ultraviolet detection. J. Chromatogr. B. Biomed. Sci. App. 700, 31–47. (doi:10.1016/s0378-4347(97)00303-4)

43. Söderlund MB, Sjöberg A, Svärd G, Fex G, Nilsson-Ehle P. 2002 Biological variation of retinoids in man. Scand. J. Clin. Lab. Invest. 62, 511–519. (doi:10.1080/003655102321004521)

44. Saltelli A, Tarantola S, Chan KP-S. 1999 A Quantitative Model-Independent Method for Global Sensitivity Analysis of Model Output. Technometrics 41, 39–56. (doi:10.1080/00401706.1999.10485594)

45. Bill A. 1964 The Albumin Exchange in the Rabbit Eye. Acta Physiol. Scand. 60, 18–29. (doi:10.1111/j.1748-1716.1964.tb02865.x)

46. Satre MA, Penner JD, Kochhar DM. 1989 Pharmacokinetic assessment of teratologically effective concentrations of an endogenous retinoic acid metabolite. Teratology 39, 341–348. (doi:10.1002/tera.1420390406)

47. Idres N, Marill J, Flexor MA, Chabot GG. 2002 Activation of retinoic acid receptor-dependent transcription by all-trans-retinoic acid metabolites and isomers. J. Biol. Chem. 277, 31491–31498. (doi:10.1074/jbc.M205016200)

48. Majumdar A, Petrescu AD, Xiong Y, Noy N. 2011 Nuclear Translocation of Cellular Retinoic Acid-binding Protein II Is Regulated by Retinoic Acid-controlled SUMOylation *. J. Biol. Chem. 286, 42749–42757. (doi:10.1074/jbc.M111.293464)

49. Dong D, Ruuska SE, Levinthal DJ, Noy N. 1999 Distinct Roles for Cellular Retinoic Acid-binding Proteins I and II in Regulating Signaling by Retinoic Acid *. J. Biol. Chem. 274, 23695–23698. (doi:10.1074/jbc.274.34.23695)

