## Supplemental material for "All-*trans* retinoic acid and fluid transport in myopigenesis"

For submission to *Royal Society Interface*

### S1. Further details of the mathematical model

#### S1.1 atRA degradation

We here provide more detail on our model of atRA degradation. atRA degradation is a complex process that is primarily mediated by CYP26A1 and CYP26B1 in mammalian cells [1–5], and which therefore depends on the concentrations of these two enzymes in cells. Importantly, the CYP enzymes can be induced by atRA, i.e. more atRA leads to greater concentrations of CYP. Jing et al. [6] have summarized the data of Thatcher et al. [7] and Tay et al. [3], and proposed that CYP26A1 induction can be described by

$$CYP26A1_{ind} = \left(\frac{Ind_{max}/f_{u,inc}}{Ind_{C50} + c_{3,S}/f_{u,inc}}\right) c_{3,S}, \qquad (S1)$$

where $CYP26A1_{ind}$ is the fold-induction of CYP26A1 above baseline levels; $Ind_{max}$ , $Ind_{C50}$ and $f_{u,inc}$ are constants taken from [6]; and $c_{3.S}$ is the atRA:SA concentration in the sclera. The term $f_{u,inc}$, the so-called "fraction unbound in the incubation", accounts for binding of atRA to materials during the *in vitro* assay used to obtain $Ind_{max}$ and $Ind_{C50}$, as summarized in [8]. Thus, the concentration of CYP26A1 in a given tissue can be written as $[CYP26A1] = CYP26A1_{ind}\ [CYP26A1]_0$, where $[CYP26A1]_0$ is the concentration of CYP26A1 in the same tissue at baseline levels of atRA. Unfortunately, we are not aware of similar induction data for CYP26B1.

In conjunction with an expression for CYP induction by atRA, we must also describe the degradation kinetics of atRA in terms of the concentration of atRA and the activity of CYP26A1 and CYP26B1. Jing et al. [6] describe the degradation via Michaelis-Menten kinetics to write the degradation rate of atRA by CYP26A1 as

$$-\frac{V_{max}[CYP26A1]}{K_m + c_{3,S}/f_{u,inc}}\,\frac{c_{3,S}}{f_{u,mic}} = -\frac{V_{max}[CYP26A1]_0}{K_m + \dfrac{c_{3,S}}{f_{u,mic}}}\left(\frac{Ind_{max}/f_{u,inc}}{Ind_{C50} + c_{3,S}/f_{u,inc}}\right)\frac{c_{3,S}^2}{f_{u,mic}}. \qquad (S2)$$

Here $V_{max}$ and $K_m$ are standard Michaelis-Menten parameters and $f_{u,mic}$ accounts for atRA availability in the micelles used to derive the Michaelis-Menten parameters. In this formulation, we would need to add a similar contribution to equation $(S2)$ to account for atRA degradation by CYP26B1. This approach suffers from needing to know a large number of parameters, each of which has its own level of uncertainty. We note that this formulation neglects the kinetics of CYP26 induction, consistent with our steady state approach.

A more feasible approach was proposed by White et al. [9], who essentially assumed that the cytoplasmic uptake of atRA was proportional to the extracellular concentration of atRA:SA. White et al. further postulated that the concentration of the CYP enzymes was given by an expression that reduces to that of equation ($S1$), albeit with different nomenclature,[1] and that the cytoplasmic degradation rate of atRA was

[1] White et al. considered the effects of spatial location and allowed the "strength" of the atRA signal to vary nonlinearly with atRA concentration. To reduce their formulation to that of equation (S2) we consider a fixed location and assume that atRA signal "strength" is linearly proportional to atRA concentration.

linearly proportional to the cytoplasmic concentration of atRA and the CYP enzymes (first order kinetics). At steady state, this formulation reduces to, using our nomenclature

$$r_{3,S} = -k_{CYPdeg}\left(\frac{Ind_{max}\ c_{3,S}/f_{u,inc}}{Ind_{C50} + c_{3,S}/f_{u,inc}}\right)c_{3,S}, \qquad (S3)$$

where $k_{CYPdeg}$ is an effective degradation rate constant that incorporates the effects of all CYP enzymes (units: per time per unit tissue volume). This approach involves fewer unknown parameters than equation (S2), but is somewhat less biologically fidelic. In particular, it assumes that the induction kinetics of all CYP enzymes are similar to those of CYP26A1, or equivalently, that atRA degradation is dominated by CYP26A1. Further, as for the first approach, it neglects the kinetics of CYP26 induction, consistent with our steady state approach. To maintain tractability of this problem, we have adopted the second approach *i.e.* equation ($S3$).

**S1.2 Summary of the equations**

In this section we summarize the final set of equations we derived the main text. The equations for the pressure $p_M$ and the concentration of solute $i = 2,3$, $c_{i,M}$ , in region $M = C, S$ are given by

$$\frac{d^2\,p_M}{dx^2} = q_M, \qquad (S4)$$

$$\frac{d}{dx}\left(-\frac{K_M}{\mu}\frac{dp_M}{dx}\,c_{i,M}\right) - \mathcal{D}_{i,M}\,\frac{d^2 c_{i,M}}{dx^2} = r_{i,M}(x) + b_{i,M}(x). \qquad (S5)$$

Here $\mathcal{D}_{i,M}$ is the diffusion coefficient of solute $i$ in region $M$, $K_M$ is the Darcy permeability in region $M$ and $\mu$ is the dynamic viscosity of extravascular fluid. The source terms in the two regions are given in the table below.

| Choroid | Sclera |
|---|---|
| $q_C = a(x)\,\mathcal{L}_p^*\left[(p_B - p_C(x)) - \sum_{i=2}^{3}\sigma_i RT\left(c_{i,B} - c_{i,C}(x)\right)\right]$ | $q_S = 0$ |
| $b_{i,C} = \beta\left(c_{i,B} - c_{i,C}\right)a(x)$ | $b_{i,S} = 0$ |
| $r_{2,C} = -k_{prod}a(x)$ | $r_{2,S} = k_{CYPdeg}\left(\frac{Ind_{max}\ c_{3,S}/f_{u,inc}}{Ind_{C50} + c_{3,S}/f_{u,inc}}\right)c_{3,S}$ |
| $r_{3,C} = k_{prod}a(x)$ | $r_{3,S} = -k_{CYPdeg}\left(\frac{Ind_{max}\ c_{3,S}/f_{u,inc}}{Ind_{C50} + c_{3,S}/f_{u,inc}}\right)c_{3,S}$ |

The boundary conditions are given by equations (11-14) and (21-24) of the main text. The equations ($S4$-$S5$) result in 6 equations for 6 unknowns: $p_M$ and $c_{i,M}$ ($i = 2, 3$) for $M = C, S$. An additional 7$^{th}$ unknown in the model is $IOP$, which is found from the algebraic equation (1): $Q_{prod} - Q_u(IOP) = \mathbb{C}(IOP - EVP)$. This equation is coupled to the main system through boundary condition (13), and pressures in the blood (equation (6)) and the SCS (equation (7)).

#### S1.3 Numerical solution

The equations for fluid transport and solute transport are two second-order, coupled, nonlinear ordinary differential equations (ODEs), defined on two domains: the choroid $0 \leq x < L_C$ and the sclera $L_C \leq x \leq L_C + L_S$. These equations were rewritten as a system of four first-order ODEs and were solved in MATLAB R2024a (The Mathworks, Natick, MA) using the boundary value solver `bvp4c`, which is fourth-order accurate and iteratively refines the mesh in the regions of the domain where the error is larger than desired.

After solving the governing equations, we extracted a number of high-level outputs, chosen because they had physiologic meaning and had been previously measured. Specifically, we extracted values for the following quantities:

- $IOP$.
- The fraction of total outflow due to unconventional outflow, $Q_u/Q_{prod}$.
- The mean SA plus atRA:SA concentration in the sclera as a percentage of blood SA concentration, $< c_2+c_3 >_S/c_{2,B}$, which to an excellent approximation (in non feeding condition) is simply $< c_2 >_S/c_{2,B}$. Here and throughout the brackets $<\cdot>$ represent a spatial averaging operation, with the subscript following the bracket indicating the domain over which the averaging is occurring, i.e. $< c_2 >_C$, $< c_2 >_S$ and $< c_2 >_{CS}$ denote the spatially averaged concentration of SA in the choroid, sclera and choroid + sclera, respectively.
- The net volumetric flow rate of fluid out of the choroidal vasculature per unit area. We computed $u_{CC} = \int_0^{L_C} q_C dx$ (m/s), where $q_C$ is defined in Eq. (9) and the product of $u_{CC}$ with the cross-sectional area of the choroidal-scleral interface is the net flow rate at which fluid leaves the vessels of the CC.
- Fluid outflow rate across the sclera per unit area, $u_S$ (m/s). The product of $u_S$ and the cross-sectional area of the choroidal-scleral interface is the net flow rate of fluid crossing the sclera.
- The net leakage rate of SA plus atRA:SA from the choroidal vasculature (mol/s), given by $A_S \int_0^{L_C} a(x)\, \beta(c_{3,B} - c_{3,C}) dx$.
- The concentration of atRA plus atRA:SA averaged over the choroid and sclera, $< c_1 + c_3 >_{CS}$, which to a good approximation is simply $< c_3 >_{CS}$. Note that here we considered both atRA feeding and non-feeding conditions.

#### S1.4. Details of the Sensitivity Analysis

Once we had optimized the input parameter values, we carried out a global sensitivity analysis, using the Fourier amplitude sensitivity test (eFAST) as proposed by Saltelli et al. [10] and Marino et al. [11]. In this variance-based spectral method each input parameter whose sensitivity is being evaluated is sampled from a uniform distribution within its corresponding range (see below). This sampling follows a periodic curve with a certain frequency. To achieve better sampling, for each parameter $N_r$ independent curves are considered, each with a random shift and $N_s$ sampling points per curve. A total sensitivity index is then generated for each parameter, which is a measure of the importance of that parameter in determining the value of the observed model output and also accounts for interactions of the parameter with other parameters. The higher the value of the sensitivity index, the greater the parameter's impact on the model output. The total sensitivity index is computed by measuring the remaining variance of the output if all parameters but the considered one are fixed.

Our sensitivity analysis followed a two-step approach similar to that described above for optimizing input parameters. First, we considered only fluid and SA transport, varying 6 relevant model input parameters:

$K_S, \mathcal{L}_p^*, \beta, u_{RPE}, \Delta P_{blood}$ and $\Delta P_{SCS}$. The parameters $K_S$ and $u_{RPE}$ were varied over the range $[0.5,1.5]$ times their reference values (table 1). The values of $\mathcal{L}_p^*$ and $\beta$ were obtained by optimization, and we therefore judged them to be less certain; we thus considered larger ranges, namely [1/3,3] times their reference (optimized) values. Finally, we chose ranges for pressures differences in the blood and the SCS to be $\Delta P_{blood} \in [3,7]$ mmHg and $\Delta P_{SCS} \in [0,2]$ mmHg. A dummy parameter was also included in the sensitivity analysis, which is an artificially introduced parameter that does not affect the model outputs and thus acts as a "negative control" for the analysis. Here we set $N_r = 3$ and $N_s = 1000$.

For this stage of the sensitivity analysis, we evaluated input parameter effects on the following model outputs: $IOP$, fluid flux out of the choroidal vessels $u_{CC}$, fluid flux through the sclera $u_S$, and spatially averaged SA concentration over the choroid and sclera $< c_2 >_{CS}$. Note that due to limited availability of experimental data, we carried out input parameter optimization using the mean SA concentration in only the sclera $< c_2 >_S$, while for the sensitivity analysis we used the mean SA concentration over both the choroid + sclera $< c_2 >_{CS}$ since that is arguably of greater physiological interest.

After the first stage of the sensitivity analysis, we then performed a second sensitivity analysis examining atRA concentrations, in which we considered all the parameters and ranges above, as well as: atRA production rate $k_{prod}$, atRA degradation rate $k_{CYPdeg}$, $Ind_{C50}$ (see equation (19)) and concentration of atRA:SA in blood $c_{3,B}$. We chose to vary only the latter two parameters in the degradation reaction equation, since it is possible to rewrite equation (18) as $r_{3,S} = -k_{CYPdeg} Ind_{max} \left( \frac{c_{3,S}}{f_{u,inc} Ind_{C50} + c_{3,S}} \right) c_{3,S}$, which shows that varying $k_{CYPdeg}$ while keeping $Ind_{max}$ constant is equivalent to varying the product $k_{CYPdeg} Ind_{max}$. Similarly, varying $Ind_{C50}$ captures the variation of the product $f_{u,inc} Ind_{C50}$. Following the reasoning in the first part of the sensitivity analysis, the most uncertain parameters $k_{prod}$ and $k_{CYPdeg}$ were varied over the range [1/3,3] times their reference (optimized) values, while $Ind_{C50}$ was varied over the range $[0.5,1.5]$ times its reference value. Here we set $N_r = 3$ and $N_s = 1500$.

### S2. Selection of parameter values

#### S2.1. Scaling of parameters between species

*Scaling of parameters between mice and humans:* We wish to use measurements of atRA concentrations obtained from mice (conditions 1 and 2 in the main text) to make inferences about the situation in humans (condition 3). Similarly, some parameter values are well-known in humans but less well-known in mice. Both situations require us to transfer parameter values between these two species, a process that is made complex by known differences in cellular metabolic rate [12] and eye size [13] between mice and humans.

Let us first consider parameter $\beta$, which is the caveolae-mediated transmural transport of SA in the choriocapillaris. Savage et al. [12] have shown that for metabolically active cells, including endothelial cells and fibroblasts, cell volume (and hence cell mass) are independent of body mass, while cellular metabolic rate, that is the rate of cellular processes, varies with body mass to the ¼ power. This scaling is a consequence of the well-known dependence of organismal metabolic rate on body mass to the ¾ power; expressed in a different way, the time scale for cellular processes varies with body mass to the -¼ power for metabolically active cells. We assume this to apply to the transmural transport of SA, characterized by the parameter $\beta$. We expect that $\beta$ will be inversely proportional to the time scale of caveolar shuttling across the endothelial cells of the choriocapillaris and will vary with body mass to the ¼ power. With this scaling approach, and using mouse and human body masses of 0.02 and 70 kg, respectively [14], we expect that $\beta$ for the mouse will be 7.7-fold higher than $\beta$ for the human.

Similar arguments and scaling hold for other active processes, like synthesis and degradation of atRA, $k_{prod}$ and $k_{CYPdeg}$. Further, we expect that $u_{RPE}$, the pumping velocity of the RPE, will also vary with body mass to the ¼ power, since it is metabolically driven. We note in passing that these scaling arguments provide general guidance but that, depending on specific inter-species details, the 7.7-fold value may not be exactly correct. However, we expect the trend to be correct, i.e. $\beta$, $k_{prod}$, $k_{CYPdeg}$ and $u_{RPE}$ should all be greater in the mouse that in the human. Indeed, through optimized fitting of the values of $\beta$ for mice and humans (see below), we actually found a significantly larger mouse-to-human ratio than predicted by scaling (approximately 404 instead of 7.7; see table 1 in main text), which highlights the limitations of the scaling approach. Conversely, we note that in our model the optimized (fitted) value of the modified capillary permeability $\mathcal{L}_p^*$ does not adhere to this scaling pattern. Indeed, the fitted value of $\mathcal{L}_p^*$ is 2 orders of magnitude higher in mouse than in human (table 1), while we do not expect any metabolic or body size dependency in $\mathcal{L}_p^*$. This highlights the limitations of this scaling approach and motivates future species-specific measurements of these quantities, as discussed in the limitations section in the main text.

##### S2.2 Fitting of the parameter values

To specify the baseline values of the uncertain parameters, we performed optimization, with the goal of reproducing several known outputs for which we judged that reasonable experimental data existed. This was undertaken in a two-step process: first we considered only fluid and SA transport, for which the uncertain input parameters were $\beta$ and $\mathcal{L}_p^*$, representing the SA leakage rate from vessels and fluid exchange rate from vessels. For this step, we attempted to match the model outputs with measured values of: IOP, the fraction of total aqueous outflow due to unconventional outflow, and mean concentration of SA in the sclera. To do so, we minimized the following cost function

$$cost = \left(\frac{IOP - IOP_t}{IOP_t}\right)^2 + \left(\frac{Q_u - Q_{u,t}}{Q_{u,t}}\right)^2 + \left(\frac{< c_2 >_S - < c_2 >_{S,t}}{< c_2 >_{S,t}}\right)^2,$$

separately in mouse and humans. In the above expression subscript $t$ refers to the target values reported in the last part of table 1. The optimization was performed using the Matlab function `fmincon`. The resulting optimized values of the input parameters are reported in table 1.

In the second step, we considered atRA transport, for which the uncertain input parameters were $k_{CYPdeg}$ and $k_{prod}$. Here we performed two separate optimizations with the goals of minimizing least square relative error of the model predictions compared to experimentally measured mean concentrations of atRA:SA in the choroid + sclera in mouse, $< c_3 >_{CS}$. First, we found the optimum value of $k_{CYPdeg}$ considering feeding conditions only. We then optimized the input value of $k_{prod}$ considering the physiological case of no feeding. This approach was convenient since the amount of atRA delivered to ocular tissues during feeding vastly overwhelms the local production, so that in the feeding situation $k_{prod}$ is relatively unimportant and the atRA concentration in the choroid + sclera is essentially determined only by $k_{CYPdeg}$. The values for human were then obtained using scaling argument described in §S2.1.

##### S2.3. Choice of parameter values in table 1

In the table below we describe the reasons behind the choices of parameters in table 1, with appropriate references. The order of parameters is the same as in table 1.

| Parameter | Explanation |
|---|---|
| $L_S$ | In mice: based on measurements of 40-100 µm in wild-type C57BL/6 mice [15]; of 36-65 µm in wild-type C57BL/6 mice (2 months old, inflated eyes) [16]; and of 41-61 µm in Wild-type C57BL/6 mice, 2-4 months old [17]. In humans: based on micro-MRI imaging of freshly enucleated human eyes [18]. |
| $L_C$ | Choroidal thickness differs significantly between humans and mice. In humans, the most reliable measurements of total choroidal thickness are made using OCT, and show that choroidal thickness depends strongly on position, refractive status, disease state and age. A map of choroidal thickness based on OCT measurements from [19] has been constructed and used in [20] to compute a mean thickness of 268 $\mu$m. We use this value. In mice, the choroid is thinner and OCT measurements do not seem to exist in the literature. Thus, we relied on histologic slices and, in particular, we used Figure 3D in [21]. Comparing the thickness of the choroid estimated from the image with the scale bar in the figure, we obtained an average thickness of the choroid of 30 $\mu$m. |
| $\alpha$ | In humans. The thickness of the choriocapillaris is not well studied. One review article [22] states that the capillaries of the choriocapillaris are 40-60 $\mu$m in diameter, while a second article [23] reports a thickness of the choriocapillaris of up to 10 $\mu$m. Curcio et al. [24] measured a thickness of 10-20 $\mu$m. In this work we used a value between the reported extremes of 30 $\mu$m, which is 1/9$^{th}$ of the total choroidal thickness. In mice, we again used Figure 3D in [21] and estimated a thickness of approximately half of that of the choroid. Following the above discussion, we thus set $\alpha$ equal to 1/9 for the human and to 1/2 for the mouse. |
| $\mu$ | We assumed that interstitial fluid has a viscosity equal to saline (water). |
| $u_{RPE}$ | In humans, fluid velocity in postoperative eyes with retinal detachment due to the RPE was measured and converted to a velocity of $3 \pm 1.5 \times 10^{-8}$ m/s [25] while cultured fetal RPE cells pumped at a velocity of $0.7 - 2.4 \times 10^{-8}$ m/s [26]. A comparable value of $3.9 \times 10^{-8}$ m/s was found in cynomolgus monkeys with retinal detachment [27]. We use $3 \times 10^{-8}$ m/s as our value. In mice, there is no data available to our knowledge. Instead, we use the scaling argument described in §S2.1 to obtain $23 \times 10^{-8}$ m/s. |
| $p_{orbit}$ | Pressure in the orbit has been estimated in a number of studies, reviewed in [28]. In human eyes, measurements lie in the range of 3-6 mmHg. There are no data available for mouse that we are aware of. We use 3 mmHg for both human and mouse. |
| $\Delta P_{blood}$ | Mäepea *et al.* [29] used a micropuncture technique to measure blood pressure in the choroidal veins and choriocapillaris of rabbits. IOP was increased stepwise to determine pressure variations in blood vessels in response to IOP changes. The author found that the pressure in the choriocapillaris was 7.6±0.5 mmHg greater than IOP. In this work we chose a value of 5 mmHg for mice and humans, and investigated the effect this parameter value in our sensitivity analysis. |
| $\Delta P_{SCS}$ | In monkeys, Emi et al. [30] measured a pressure difference of 0.8 to 3.7 mmHg depending on position, which is consistent with the concept that the quantity $\Delta P_{scs}$ can be interpreted as the pressure drop along the unconventional pathway between the anterior chamber and the measurement location. Since our computational domain is at an "average" location within the eye, we prescribe a constant, intermediate value. |
| $EVP$ | In mice: EVP was measured to be 5.4 mmHg in male BALB/cJ mice 30 to 42 weeks of age [31] and 9.6 mmHg in NIH Swiss white mice of age 8-12 weeks [32]. We took 7.5 mmHg as an average. In humans: In a cohort of human volunteers, daytime EVP varied between 6.4 and 7.8 mmHg, depending on posture [33]. We used an average value of 7.1 mmHg. |

| | |
|---|---|
| $Q_{prod}$ | In mice: Based on updated calculations described in [34] accounting for corrected posterior chamber volume in mice. In humans: Average data from Table 1 of [35] for normal humans. |
| $\mathbb{C}$ | In mice: Taken from measurements of enucleated eyes from 66 10–14 week old male C57BL/6J mice [36]. In humans: Based on bilateral tonographic measurements in 34 healthy human volunteers (average of daytime and nocturnal values) [37]. |
| $\mathcal{L}_p^*$ | Value obtained by optimization, see section §S2.2. |
| $K_S$ | In mice: Figure 7(b) from [15] gives $K_S = 4.3 \times 10^{-17}$m$^2$ for wild-type C57BL/6 mice at 37 °C. Note that this is the value extrapolated to zero compressive strain, and that there is significant spread in this value (almost one order of magnitude). Due to the nature of the experiments, the measured value is primarily for flow parallel to the collagen fibers in the sclera, while our model considered flow in the direction normal to this. Thus, the measured value should be considered as an upper bound. Here we use $K_S = 6.4 \times 10^{-18}$ m$^2$. In humans: We use the data from Fatt and Hedbys [38] that, assuming a fluid viscosity of 0.7 × $10^{-3}$ Pa·s, gives a value of 1.33 × $10^{-18}$ m$^2$. Another measurement is also available by Jackson *et al.* [39] of 5.85 × $10^{-18}$ m$^2$. We test the dependency of this parameter in our sensitivity analysis. |
| $K_C$ | The choroid cannot be easily separated from the RPE and Bruch's membrane; thus the existing studies on permeability to water refer to such layers together. Tsuboi [40] measured the hydraulic conductivity of the RPE-choroid complex and found a value of 1.6 × $10^{-11}$ m/Pa/s. Moore et al. [41] found that the hydraulic conductivity of the Bruch's membrane-choroid complex varies significantly with age, from ~$10^{-8}$ m/Pa/s in youth to ~$10^{-10}$ m/Pa/s in elderly. Ruffini et al. [42] made use of a mathematical model to speculate that the value proposed by Tsuboi's is likely too small. Assuming a thickness of the choroid of 268 µm and a viscosity of the fluid of 0.7 × $10^{-3}$ Pa·s, Moore's measurements imply that the permeability of the Bruch's membrane choroid complex (treated as a homogeneous tissue) ranges between ~2 × $10^{-17}$ m$^2$ and ~2 × $10^{-15}$ m$^2$. Noting that most of the resistance to water flow across these layers is likely due to Bruch's membrane, it follows that the permeability of the choroid is much smaller than that of the sclera. In this work we set $K_C = 100\ K_S$ for both mice and humans. |
| $\mathcal{D}_{2,S}$, $\mathcal{D}_{3,S}$ | Due to the small molecular weight of atRA compared to SA, the binding of atRA to SA is assumed to not affect the diffusivity of SA, so that $\mathcal{D}_{2,S} = \mathcal{D}_{3,S}$. A number of authors have measured diffusivity of albumin in sclera, finding a range of values that appear to be species-dependent. In human, tabulated values in [43] range from $0.61$ to $1.5 \times\ 10^{-12}$ m$^2$/s. Anderson et al.'s [44] measurements, when converted to a diffusivity assuming a typical scleral thickness of 670 um (see note 14), give $9.6 \times\ 10^{-12}$ m$^2$/s. Ambati et al. [45] summarize values for bovine in the range of $0.89$ to $2.0 \times\ 10^{-12}$ m$^2$/s, and a value for rabbit of $13 \times\ 10^{-12}$ m$^2$/s. The rabbit value is extremely large and may be an outlier. Even eliminating the rabbit data, there is a large range; we have chosen to use the value given by Anderson et al. We note that, fortunately, the model's sensitivity to this value is small. Mouse: We are unaware of any mouse-specific data and thus have assumed that the diffusivity in mouse sclera is similar to human. Note that the diffusivity of serum albumin in free solution is $875 \times\ 10^{-13}$ m$^2$/s, indicating significant hindrance by the sclera. (Free solution diffusivity from [46] in saline at pH 7, corrected to body temperature.) |
| $\mathcal{D}_{2,C}$, $\mathcal{D}_{3,C}$ | We assume that diffusivity in the choroid is the same as that in free solution due to the porous structure of the choroid vs. the sclera. This is possibly an overestimate, but the results of the model are insensitive to the assumed value. Similar to the situation in the sclera, we assume that $\mathcal{D}_{2,C} = \mathcal{D}_{3,C}$. |

| | |
|---|---|
| $\beta$ | Value obtained by optimization, see section §S2.2. |
| $c_{2,B} + c_{3,B}$ | In mice: We used data from [47] for C57BL/6 mice of both sexes, 8-10 weeks of age and converted to a molar concentration using a molecular weight of 66.5 kDa. In humans: We used the midpoint of the commonly accepted reference range for humans (42.5 g/L [48]) and converted to a molar concentration using a molecular weight of 66.5 kDa. |
| $c_{3,B}$ | In mice: We made direct measurements under atRA feeding and control conditions, as described and reported in the text. In humans: Normal atRA levels are 2.5 nM [49]. |
| $k_{prod}$ | In mice: optimized, see section §S2.2. In humans: The values for human were obtained by scaling the value for the mouse, using the scaling argument described in section §S2.1.<br>We also benchmarked the value against data from Mertz and Wallman [50], who measured an atRA production rate in chick choroid of approximately 0.73 pmol/hr in 8 mm diameter choroidal punches (control eyes). Assuming a chick choroidal thickness of 60 microns, the volume of such a punch is $3 \times 10^{-9}$ $m^3$, yielding an atRA synthesis rate of $6.7\times 10^{-8}$ $mol/m^3/s$, which is the same order of magnitude as our optimized values. We note, however, that atRA appears to play a different role in myopigenesis in chicks vs. mammals [51], and thus it is not clear how reliable the extrapolation of the value from [50] to mammals is. |
| $k_{CYPdeg}$ | In mice: optimized, see section §S2.2. In humans: The values for human were obtained by scaling the value for the mouse, using the scaling argument described in section §S2.1. |
| $Ind_{max}$ | Maximum fold induction of atRA degradation, taken directly from Jing et al. [6] |
| $Ind_{C50}$ | atRA concentration at which 50% of the maximum degradation induction occurs, taken directly from Jing et al. [6] |
| $f_{u,inc}$ | Fraction unbound in the incubation, taken directly from Jing et al. [6] |
| **Optimization targets** | |
| $IOP$ | In mice: It is known that age, time of day, and extent of anesthesia affects IOP, and that tonometry can misestimate IOP [52], i.e. there is no single, “true” value of IOP. John et al. measured mean IOP of 12.3 mmHg in anesthetized C57BL/6J mice by direct cannulation [52], a value that was later updated to 13.4 mmHg during the daytime and approximately 16 mmHg at night [53]. Lathrop et al. measured an IOP of 14.5 ± 2.6 mmHg (Mean ±SD) in a large cohort of C57BL/6 mice by tonometry [54]. We chose a value of 14.4 mmHg lying within this range. In humans: Many studies have measured IOP in large human populations, showing a dependence on age, race and sex. For example, Hollows and Graham [55] report a IOP of 15.9 ± 2.9 mmHg in 1,873 mostly Caucasian males, while Liu et al. [56] report mean values of 15.4 (95% CI: 9.1-21.6) mmHg for men and 14.9 (95% CI: 9.0-20.8) mmHg for women in a Chinese population, all by tonometry. We chose a value of 14.9 mmHg within these and other reported ranges. |
| $Q_u/Q_{\text{prod}}$ | In humans: measured values show a wide range, and we chose a value slightly higher than the percentage measured by direct tracer experiments in living human eyes but lower than the average value obtained by indirect approaches, as shown in Table 4 of [57]. In mice: the values are even less well known [57]; thus, we have assumed mouse unconventional outflow faction to be the same as in human eyes. |
| $\frac{< c_2 >_S}{c_{2,B}}$ | Allansmith et al. [58] measured albumin concentrations in the sclera and choroid of post mortem human eyes, but suffered from an extremely large range of values (suggesting unreliability) and failure to wash blood out of tissue before the assay. Extravascular albumin concentrations in the |

| | |
|---|---|
| | choroid have also been measured in monkey by Toris et al. [59], but again show a large spread. We thus did not use choroidal values, and instead simply used the mean value for albumin concentration in the sclera from Allansmith et al. [58]. |
| $< c_3 >_{cs}$ | In mice: We used values of atRA concentration in RPE + choroid + sclera measured experimentally under atRA feeding and non-feeding conditions [60]. In humans: We did not carry out optimization using this quantity in humans. |

**Table S1:** Rationale for choices of parameter values listed in Table 1 of the main text.

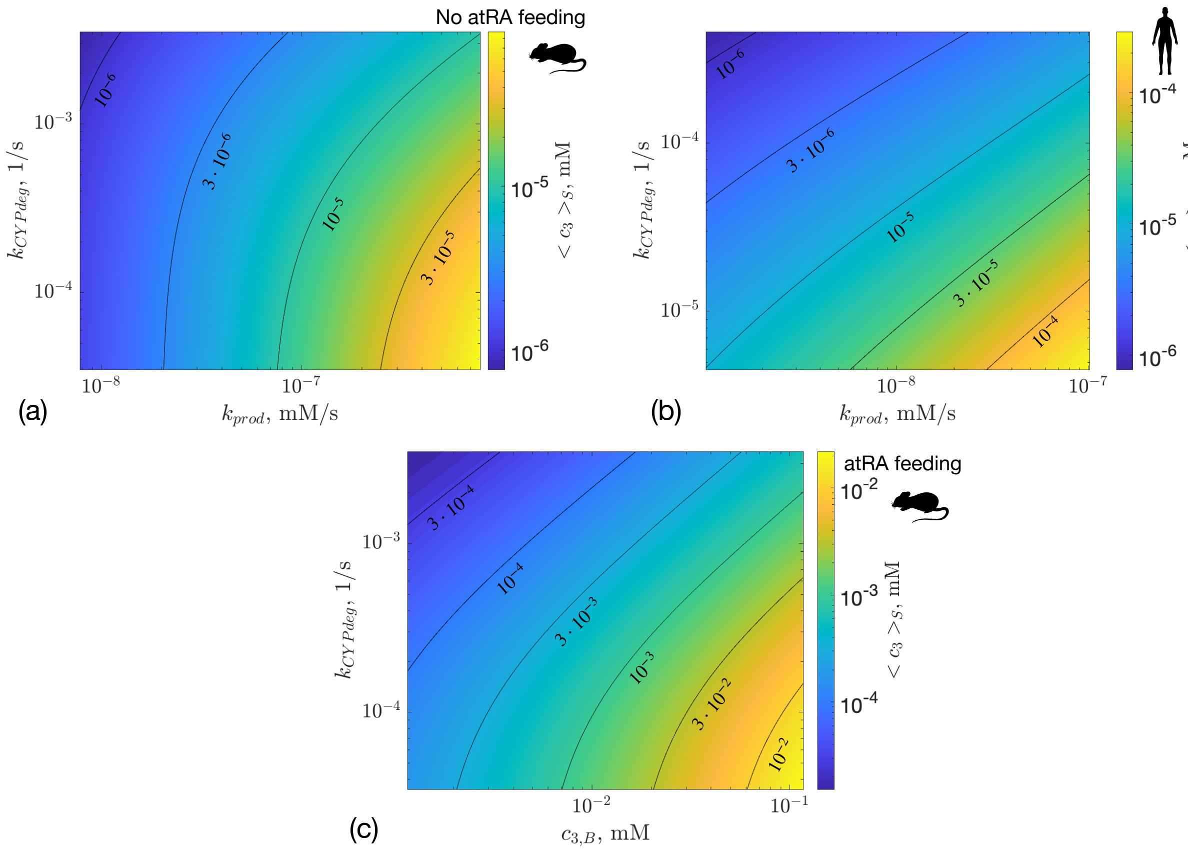


**Figure S1:** Mean atRA concentration in the sclera $< c_3 >_S$ as a function of key model parameters. Panels (a) and (b) show the dependency of $< c_3 >_S$ on the atRA synthesis rate $k_{prod}$ and atRA degradation rate $k_{CYPdeg}$ for (a) mouse and (b) human. In panel (c) we consider the atRA feeding scenario and show the dependency of $< c_3 >_S$ on atRA concentration in the blood plasma $c_{3,B}$ and $k_{CYPdeg}$. The black curves show isoconcentration contours with adjacent concentration values displayed.

#### References

1. Topletz AR, Tripathy S, Foti RS, Shimshoni JA, Nelson WL, Isoherranen N. 2015 Induction of CYP26A1 by Metabolites of Retinoic Acid: Evidence That CYP26A1 Is an Important Enzyme in the Elimination of Active Retinoids. *Mol. Pharmacol.* **87**, 430–441. (doi:10.1124/mol.114.096784)

2. Isoherranen N, Zhong G. 2019 Biochemical and physiological importance of the CYP26 retinoic acid hydroxylases. *Pharmacol. Ther.* **204**, 107400. (doi:10.1016/j.pharmthera.2019.107400)

3. Tay S, Dickmann L, Dixit V, Isoherranen N. 2010 A Comparison of the Roles of Peroxisome Proliferator-Activated Receptor and Retinoic Acid Receptor on CYP26 Regulation. *Mol. Pharmacol.* **77**, 218–227. (doi:10.1124/mol.109.059071)

4. Topletz AR, Thatcher JE, Zelter A, Lutz JD, Tay S, Nelson WL, Isoherranen N. 2012 Comparison of the function and expression of CYP26A1 and CYP26B1, the two retinoic acid hydroxylases. *Biochem. Pharmacol.* **83**, 149–163. (doi:10.1016/j.bcp.2011.10.007)

5. Ghaffari H, Petzold LR. 2018 Identification of influential proteins in the classical retinoic acid signaling pathway. *Theor. Biol. Med. Model.* **15**, 16. (doi:10.1186/s12976-018-0088-7)

6. Jing J, Nelson C, Paik J, Shirasaka Y, Amory JK, Isoherranen N. 2017 Physiologically Based Pharmacokinetic Model of All-trans-Retinoic Acid with Application to Cancer Populations and Drug Interactions. *J. Pharmacol. Exp. Ther.* **361**, 246–258. (doi:10.1124/jpet.117.240523)

7. Thatcher JE, Zelter A, Isoherranen N. 2010 The relative importance of CYP26A1 in hepatic clearance of all-trans retinoic acid. *Biochem. Pharmacol.* **80**, 903–912. (doi:10.1016/j.bcp.2010.05.023)

8. Bowman CM, Benet LZ. 2018 An examination of protein binding and protein-facilitated uptake relating to *in vitro-in vivo* extrapolation. *Eur. J. Pharm. Sci.* **123**, 502–514. (doi:10.1016/j.ejps.2018.08.008)

9. White RJ, Nie Q, Lander AD, Schilling TF. 2007 Complex regulation of cyp26a1 creates a robust retinoic acid gradient in the zebrafish embryo. *PLoS Biol.* **5**, e304. (doi:10.1371/journal.pbio.0050304)

10. Saltelli A, Tarantola S, Chan KP-S. 1999 A Quantitative Model-Independent Method for Global Sensitivity Analysis of Model Output. *Technometrics* **41**, 39–56. (doi:10.1080/00401706.1999.10485594)

11. Marino S, Hogue IB, Ray CJ, Kirschner DE. 2008 A methodology for performing global uncertainty and sensitivity analysis in systems biology. *J. Theor. Biol.* **254**, 178–196. (doi:10.1016/j.jtbi.2008.04.011)

12. Savage VM, Allen AP, Brown JH, Gillooly JF, Herman AB, Woodruff WH, West GB. 2007 Scaling of number, size, and metabolic rate of cells with body size in mammals. *Proc. Natl. Acad. Sci.* **104**, 4718–4723. (doi:10.1073/pnas.0611235104)

13. Howland HC, Merola S, Basarab JR. 2004 The allometry and scaling of the size of vertebrate eyes. *Vision Res.* **44**, 2043–2065. (doi:10.1016/j.visres.2004.03.023)

14. Davies B, Morris T. 1993 Physiological Parameters in Laboratory Animals and Humans. *Pharm. Res.* **10**, 1093–1095. (doi:10.1023/A:1018943613122)

15. Brown DM, Pardue MT, Ethier CR. 2021 A biphasic approach for characterizing tensile, compressive and hydraulic properties of the sclera. *J. R. Soc. Interface* **18**, 20200634. (doi:10.1098/rsif.2020.0634)

16. Myers KM, Cone FE, Quigley HA, Gelman S, Pease ME, Nguyen TD. 2010 The in vitro inflation response of mouse sclera. *Exp. Eye Res.* **91**, 866–875. (doi:S0014-4835(10)00304-0 [pii] 10.1016/j.exer.2010.09.009)

17. Cone-Kimball E, Nguyen C, Oglesby EN, Pease ME, Steinhart MR, Quigley HA. 2013 Scleral structural alterations associated with chronic experimental intraocular pressure elevation in mice. *Mol. Vis.* **19**, 2023–39.

18. Norman RE, Flanagan JG, Rausch SMK, Sigal IA, Tertinegg I, Eilaghi A, Portnoy S, Sled JG, Ethier CR. 2010 Dimensions of the human sclera: Thickness measurement and regional changes with axial length. *Exp. Eye Res.* **90**, 277–284. (doi:S0014-4835(09)00319-4 [pii] 10.1016/j.exer.2009.11.001)

19. Hoseini-Yazdi H, Vincent SJ, Collins MJ, Read SA, Alonso-Caneiro D. 2019 Wide-field choroidal thickness in myopes and emmetropes. *Sci. Rep.* **9**, 3474. (doi:10.1038/s41598-019-39653-w)

20. Tweedy JH, Dvoriashyna M, Crawshaw JR, Overby DR, Repetto R, Roberts PA, Spelman TA, Stewart PA, Foss AJE. 2024 A model of the mechanisms underpinning unconventional aqueous humor outflow. *Invest. Ophthalmol. Vis. Sci.* **submitted**.

21. Matsumoto H, Mukai R, Hoshino J, Oda M, Matsuzaki T, Ishizaki Y, Shibasaki K, Akiyama H. 2021 Choroidal congestion mouse model: Could it serve as a pachychoroid model? *PloS One* **16**, e0246115. (doi:10.1371/journal.pone.0246115)

22. Pichi F, Aggarwal K, Neri P, Salvetti P, Lembo A, Nucci P, Gemmy Cheung CM, Gupta V. 2018 Choroidal biomarkers. *Indian J. Ophthalmol.* **66**, 1716–1726. (doi:10.4103/ijo.IJO_893_18)

23. Nickla DL, Wallman J. 2010 The multifunctional choroid. *Prog. Retin. Eye Res.* **29**, 144–168. (doi:10.1016/j.preteyeres.2009.12.002)

24. Curcio CA, Messinger JD, Sloan KR, Mitra A, McGwin G, Spaide RF. 2011 Human chorioretinal layer thicknesses measured in macula-wide, high-resolution histologic sections. *Invest. Ophthalmol. Vis. Sci.* **52**, 3943–3954. (doi:10.1167/iovs.10-6377)

25. Chihara E, Nao-i N. 1985 Resorption of subretinal fluid by transepithelial flow of the retinal pigment epithelium. *Graefes Arch. Clin. Exp. Ophthalmol.* **223**, 202–204. (doi:10.1007/BF02174060)

26. Adijanto J, Banzon T, Jalickee S, Wang NS, Miller SS. 2009 CO2-induced ion and fluid transport in human retinal pigment epithelium. *J. Gen. Physiol.* **133**, 603–622. (doi:10.1085/jgp.200810169)

27. Pederson JE, Cantrill HL. 1984 Experimental Retinal Detachment: V. Fluid Movement Through the Retinal Hole. *Arch. Ophthalmol.* **102**, 136–139. (doi:10.1001/archopht.1984.01040030114048)

28. Enz TJ, Tschopp M. 2022 Assessment of Orbital Compartment Pressure: A Comprehensive Review. *Diagn. Basel Switz.* **12**, 1481. (doi:10.3390/diagnostics12061481)

29. Mäepea O. 1992 Pressures in the anterior ciliary arteries, choroidal veins and choriocapillaris. *Exp. Eye Res.* **54**, 731–736. (doi:10.1016/0014-4835(92)90028-Q)

30. Emi K, Pederson JE, Toris CB. 1989 Hydrostatic pressure of the suprachoroidal space. *Invest. Ophthalmol. Vis. Sci.* **30**, 233–238.

31. Millar JC, Clark AF, Pang I-H. 2011 Assessment of Aqueous Humor Dynamics in the Mouse by a Novel Method of Constant-Flow Infusion. *Invest. Ophthalmol. Vis. Sci.* **52**, 685–694. (doi:10.1167/iovs.10-6069)

32. Aihara M, Lindsey JD, Weinreb RN. 2003 Aqueous humor dynamics in mice. *Invest. Ophthalmol. Vis. Sci.* **44**, 5168–5173. (doi:10.1167/iovs.03-0504)

33. Arora N, McLaren JW, Hodge DO, Sit AJ. 2017 Effect of Body Position on Epsicleral Venous Pressure in Healthy Subjects. *Invest. Ophthalmol. Vis. Sci.* **58**, 5151–5156. (doi:10.1167/iovs.17-22154)

34. Kim D *et al.* 2024 In vivo quantification of anterior and posterior chamber volumes in mice: implications for aqueous humor dynamics. *BioRxiv Prepr. Serv. Biol.* , 2024.07.24.604989. (doi:10.1101/2024.07.24.604989)

35. McLaren JW. 2009 Measurement of aqueous humor flow. *Exp. Eye Res.* **88**, 641–647. (doi:10.1016/j.exer.2008.10.018)

36. Sherwood JM, Reina-Torres E, Bertrand JA, Rowe B, Overby DR. 2016 Measurement of Outflow Facility Using iPerfusion. *PLOS ONE* **11**, e0150694. (doi:10.1371/journal.pone.0150694)

37. Sit AJ, Nau CB, McLaren JW, Johnson DH, Hodge D. 2008 Circadian variation of aqueous dynamics in young healthy adults. *Invest. Ophthalmol. Vis. Sci.* **49**, 1473–1479. (doi:10.1167/iovs.07-1139)

38. Fatt I, Hedbys BO. 1970 Flow of water in the sclera. *Exp. Eye Res.* **10**, 243–249. (doi:10.1016/s0014-4835(70)80035-5)

39. Jackson TL, Hussain A, Hodgetts A, Morley AMS, Hillenkamp J, Sullivan PM, Marshall J. 2006 Human scleral hydraulic conductivity: age-related changes, topographical variation, and potential scleral outflow facility. *Invest. Ophthalmol. Vis. Sci.* **47**, 4942–4946. (doi:10.1167/iovs.06-0362)

40. Tsuboi S. 1987 Measurement of the volume flow and hydraulic conductivity across the isolated dog retinal pigment epithelium. *Invest. Ophthalmol. Vis. Sci.* **28**, 1776–1782.

41. Moore DJ, Hussain AA, Marshall J. 1995 Age-related variation in the hydraulic conductivity of Bruch's membrane. *Invest. Ophthalmol. Vis. Sci.* **36**, 1290–1297.

42. Ruffini A, Dvoriashyna M, Govetto A, Romano MR, Repetto R. 2024 A Mathematical Model of Interstitial Fluid Flow and Retinal Tissue Deformation in Macular Edema. *Invest. Ophthalmol. Vis. Sci.* **65**, 19. (doi:10.1167/iovs.65.11.19)

43. Maurice DM, Polgar J. 1977 Diffusion across the sclera. *Exp. Eye Res.* **25**, 577–582. (doi:10.1016/0014-4835(77)90136-1)

44. Anderson OA, Jackson TL, Singh JK, Hussain AA, Marshall J. 2008 Human Transscleral Albumin Permeability and the Effect of Topographical Location and Donor Age. *Invest. Ophthalmol. Vis. Sci.* **49**, 4041–4045. (doi:10.1167/iovs.07-1660)

45. Ambati J, Canakis CS, Miller JW, Gragoudas ES, Edwards A, Weissgold DJ, Kim I, Delori FC, Adamis AP. 2000 Diffusion of High Molecular Weight Compounds through Sclera. *Invest. Ophthalmol. Vis. Sci.* **41**, 1181–1185.

46. Gaigalas AK, Hubbard JB, McCurley M, Woo S. 1992 Diffusion of bovine serum albumin in aqueous solutions. *J. Phys. Chem.* **96**, 2355–2359. (doi:10.1021/j100184a063)

47. Zaias J, Mineau M, Cray C, Yoon D, Altman NH. 2009 Reference Values for Serum Proteins of Common Laboratory Rodent Strains. *J. Am. Assoc. Lab. Anim. Sci.* **48**.

48. Busher JT. 1990 Serum Albumin and Globulin. In *Clinical Methods: The History, Physical, and Laboratory Examinations* (eds HK Walker, WD Hall, JW Hurst), Boston: Butterworths.

49. Kane MA, Napoli JL. 2010 Quantification of Endogenous Retinoids. *Methods Mol. Biol. Clifton NJ* **652**, 1–54. (doi:10.1007/978-1-60327-325-1_1)

50. Mertz JR, Wallman J. 2000 Choroidal Retinoic Acid Synthesis: A Possible Mediator between Refractive Error and Compensatory Eye Growth. *Exp. Eye Res.* **70**, 519–527. (doi:10.1006/exer.1999.0813)

51. Brown DM, Mazade R, Clarkson-Townsend D, Hogan K, Datta Roy PM, Pardue MT. 2022 Candidate pathways for retina to scleral signaling in refractive eye growth. *Exp. Eye Res.* **219**, 109071. (doi:10.1016/j.exer.2022.109071)

52. John SWM, Hagaman JR, MacTaggart TE, Peng L, Smithesf O. 1997 Intraocular Pressure in Inbred Mouse Strains. *Invest. Ophthalmol. Vis. Sci.* **38**, 249–253.

53. Savinova O, Sugiyama F, Martin J, Tomarev S, Paigen B, Smith R, John SWM. 2001 Intraocular pressure in genetically distinct mice: an update and strain survey. *BMC Genet.* **2**, 12. (doi:10.1186/1471-2156-2-12)

54. Yun H, Lathrop KL, Yang E, Sun M, Kagemann L, Fu V, Stolz DB, Schuman JS, Du Y. 2014 A laser-induced mouse model with long-term intraocular pressure elevation. *PloS One* **9**, e107446. (doi:10.1371/journal.pone.0107446)

55. Hollows FC, Graham PA. 1966 Intra-ocular pressure, glaucoma, and glaucoma suspects in a defined population. *Br. J. Ophthalmol.* **50**, 570–586.

56. Liu X, Pan X, Ma Y, Jin C, Wang B, Ning Y. 2022 Variation in intraocular pressure by sex, age, and geographic location in China: A nationwide study of 284,937 adults. *Front. Endocrinol.* **13**, 949827. (doi:10.3389/fendo.2022.949827)

57. Johnson M, McLaren JW, Overby DR. 2017 Unconventional aqueous humor outflow: A review. *Exp. Eye Res.* **158**, 94–111. (doi:10.1016/j.exer.2016.01.017)

58. Allansmith MR, Whitney CR, McClellan BH, Newman LP. 1973 Immunoglobulins in the human eye. Location, type, and amount. *Arch. Ophthalmol. Chic. Ill 1960* **89**, 36–45. (doi:10.1001/archopht.1973.01000040038010)

59. Toris CB, Pederson JE, Tsuboi S, Gregerson DS, Rice TJ. 1990 Extravascular albumin concentration of the uvea. *Invest. Ophthalmol. Vis. Sci.* **31**, 43–53.

60. Brown DM, Yu J, Kumar P, Paulus QM, Kowalski MA, Patel JM, Kane MA, Ethier CR, Pardue MT. 2023 Exogenous All-Trans Retinoic Acid Induces Myopia and Alters Scleral Biomechanics in Mice. *Invest. Ophthalmol. Vis. Sci.* **64**, 22. (doi:10.1167/iovs.64.5.22)
